# A statistical framework for inferring genetic requirements from embryo-scale single-cell sequencing experiments

**DOI:** 10.1101/2025.04.03.646654

**Authors:** Madeleine Duran, Eliza Barkan, Amy Tresenrider, Heidi Lee, Ryan Z Friedman, Nicholas Lammers, Marazzano Colón, Jennifer Franks, Brent Ewing, David Kimelman, Cole Trapnell

## Abstract

Improvements in single-cell sequencing have enabled phenotyping at organism-scale and molecular resolution, but interpreting such experiments poses computational challenges. Identifying the genes and cell types directly impacted by genetic, chemical, or environmental perturbations requires explicit modeling of lineage relationships amongst many cell types, over time, from datasets with millions of cells collected from thousands of specimens. We describe two software tools, “Hooke” and “Platt”, which exploit the rich statistical patterns within single-cell datasets to characterize the direct molecular and cellular consequences of experimental perturbations. We apply Hooke and Platt to a single-cell atlas of thousands of perturbed zebrafish embryos to synthesize a coherent map of lineage dependencies and leverage it to reveal previously unappreciated roles for fate-determining transcription factors. We show that cell type covariation in single-cell datasets is a powerful source of information for inferring how cells depend on genes and one another in the program of vertebrate development.

## Introduction

Single-cell genomics technology has advanced at a blistering pace. The throughput of single-cell transcriptome sequencing has increased by four orders of magnitude in the past five years alone^1–4^. The resulting economies of scale, in conjunction with sample multiplexing techniques, now enable single-cell genomics to not only catalog cell types but to comprehensively study the effects of well-controlled perturbations on animal development or characterize the evolution of disease pathogenesis at whole-animal scale and molecular resolution. Single-cell genomics experiments that compare healthy to diseased tissue or track the development of embryos over time typically aim to identify subpopulations of cells that differ between conditions or timepoints at the molecular level. When studying how a gene contributes to embryonic development, one might follow the “trajectories” cells take through possible gene expression states as they make fate decisions and measure how these decisions are altered by perturbing the gene in question. Both settings aim to define the cell types that change across conditions and the genes that mediate those changes.

In principle, single-cell genomics could serve as an extraordinarily high-content means of phenotyping and has prompted calls for the systematic collection of vast amounts of data into an “atlas” of perturbations^5^. Trained on a sufficiently large corpus of molecular, cellular, and tissue phenotypes, a new generation of “foundational” AI models could computationally evaluate hypotheses in minutes that otherwise would require experiments that are difficult or impossible to perform in the lab^5,6^. However, the scale and complexity of such experiments poses new, daunting computational and statistical challenges^6^. Intense efforts are underway to develop software tools to develop bioinformatic and machine learning tools to analyze single-cell perturbation datasets^7,8^. Recent tools draw from a wide range of techniques, including deep learning and foundation models as well as more conventional statistical approaches including regression or Bayesian modeling, to predict mechanisms of action for drugs, anticipate effects on gene regulation of different cell types, locate novel genetic interactions, and more. For example, scGen^9^ and CPA^10^ use variational autoencoders to model changes within the transcriptome of individual cells in response to both seen and as-yet-untested genetic perturbations. Other tools, such as CellOracle^11^, SCENIC+^12^, and D-SPIN^13^, adopt a “bottom-up” approach, first learning gene regulatory networks and then using them to predict how interventions in the network will impact each cell’s transcriptome. Some tools for comparing perturbations to controls, such as Propeller^14^, perform differential analysis on the proportions of cell types within a previously defined ontology, while others such as MILO and MELD do not require cell type annotations ahead of time^15^.

Despite the explosion of algorithms, machine learning methods, and software tools for handling single-cell perturbation experiments, there remain open problems and unmet challenges, particularly in the contexts of developmental genetics and disease pathogenesis. A first challenge is that most perturbation experiments are performed in cell lines (e.g. CROP-seq^16^), and accordingly most tools are designed to model changes in the transcriptome of a single cell type that might exist in several similar molecular states. Few, if any, tools exist for analyzing effects at the scale of complex tissues, let alone whole embryos, where perturbations can alter both the proportions and molecular states of hundreds of distinct cell types. A second challenge is that effects on progenitors impact their descendants in the cell lineage, and the network of lineage and signaling dependences between cell types is thus needed to locate those most directly impacted by a perturbation. A third challenge is that tracking how a perturbation’s effects ripple through the system can be crucial for discriminating causal relationships between genes from coincidental ones, especially in the developing embryo, but few tools explicitly model time or cell lineage relationships when performing contrasts, and none do so simultaneously. Inferring new cell lineage relationships, genetic requirements, and ultimately mapping the genetic circuits that control cell fate and function from single-cell perturbation experiments requires a computational framework that meets all these challenges at once. There is therefore an urgent need for algorithms, statistical methods, and software that meets the challenges posed by single-cell analysis at “atlas scale”.

Here, we describe two new algorithms that analyze perturbation experiments performed with single-cell transcriptomics to define how cells transition between cell states and characterize the molecular changes that occur as they do so. First, we introduce Hooke, a software package that estimates how the proportions of cells in different molecular states change across specimens, over time, in response to perturbations, and across varying experimental conditions. Hooke introduces the use of Poisson-log Normal (PLN) networks^17^ for modeling how proportions of different cell types co-vary in single cell genomics experiments. We show that the correlation structure between cell abundances is a crucial source of statistical information that reveals lineage relationships between cell types and links pathological cell states to the healthy ones that give rise to them. Second, we introduce Platt, which analyzes one or more perturbation experiments to produce a map of direct transitions that cells can make between molecular states. When provided with time-series perturbation experiments that target genes or pathways required by one or more cell types, Platt follows how other cell types are subsequently depleted and is thus able to identify cells’ ancestral states. Hooke and Platt work in concert to detect transcriptional and cell abundance phenotypes and relate them to one another, which helps users understand how disruptions to gene regulation at upstream cell types or states lead to losses in cell types downstream. Although they work with the single-cell perturbation experiments now routinely used to explore the role of a gene or compare diseased specimens to healthy controls, both tools are designed to accommodate millions of cells from thousands of specimens collected across many perturbations.

We first demonstrate these tools through an extensive reanalysis of previously published single-cell perturbation experiments from a mouse model of disease pathogenesis and a zebrafish model of embryonic development. The zebrafish embryo is a well-established system for studying development that has been used extensively in forward genetic screens as well as targeted genetic, chemical, and environmental perturbation experiments. The rapid development and high fecundity of the zebrafish makes it particularly well-suited for statistically powered sequencing experiments because hundreds of “replicate” embryos can be collected at once. We use Hooke and Platt to revise and improve our map of cell types in the developing zebrafish embryo and link them together into an embryo-scale transition graph. We then analyze patterns of gene expression over this graph to identify transcription factors putatively required for each transition. Amongst the most recurrently implicated genes were *lmx1ba* and *lmx1bb*, which we found to exhibit striking patterns of activation in many cell fates. By deeply sequencing wild type embryos and those lacking *lmx1b* factors, we demonstrate that selective expression of these factors is required for the activation of a shared connective tissue program in many of these cell types.

## Results

### Hooke performs differential analysis of cellular composition in single-cell perturbation experiments

In order to locate the cell types within embryos or complex tissues that are impacted by a perturbation, we sought to devise a statistical method that could isolate changes in each cell type’s proportions while controlling for other technical effects like batch or dissociation protocol. However, modeling observed counts of different cell types as a function of biological and technical covariates is nontrivial, because the covariates may not be independent (e.g. time and sample collections are related), the count data may be sparse (e.g. because sampling many cells per sample is costly), and there may be a complex correlation structure between the cell types (e.g. progenitors and descendants are inversely correlated in their proportions). We therefore required a statistical framework that accommodates sparse count data, flexibly describes cell proportions as a function of both biological and technical covariates, and learns the covariance structure of cell proportions from the data provided to it. We identified Poisson Lognormal (PLN) models as a promising candidate framework for building our new statistical method, which we call “Hooke”, named in honor of Robert Hooke, who first described cells and coined the term^18^.

Hooke is designed around the PLNmodels package^19^ to perform differential analysis of cell state abundances in single-cell RNA-seq experiments. As **input**, Hooke accepts a matrix containing the number of cells of each type or state observed in each sample, along with metadata for each sample (e.g. the experimental treatments applied to each sample, the time it was collected, etc.) (**Figure 1A**). Users must assign cells to types or states before running Hooke via clustering, marker-based annotation, or projecting them onto a reference data set^20,21^. As **output**, Hooke produces a fitted PLN network model through which users can visualize the cell state abundance changes that occur following each experimental perturbation. Users can also query the model to understand how cell types co-vary across samples to understand how subsets of cell types are gained at the expense of others.

**Figure 1.**
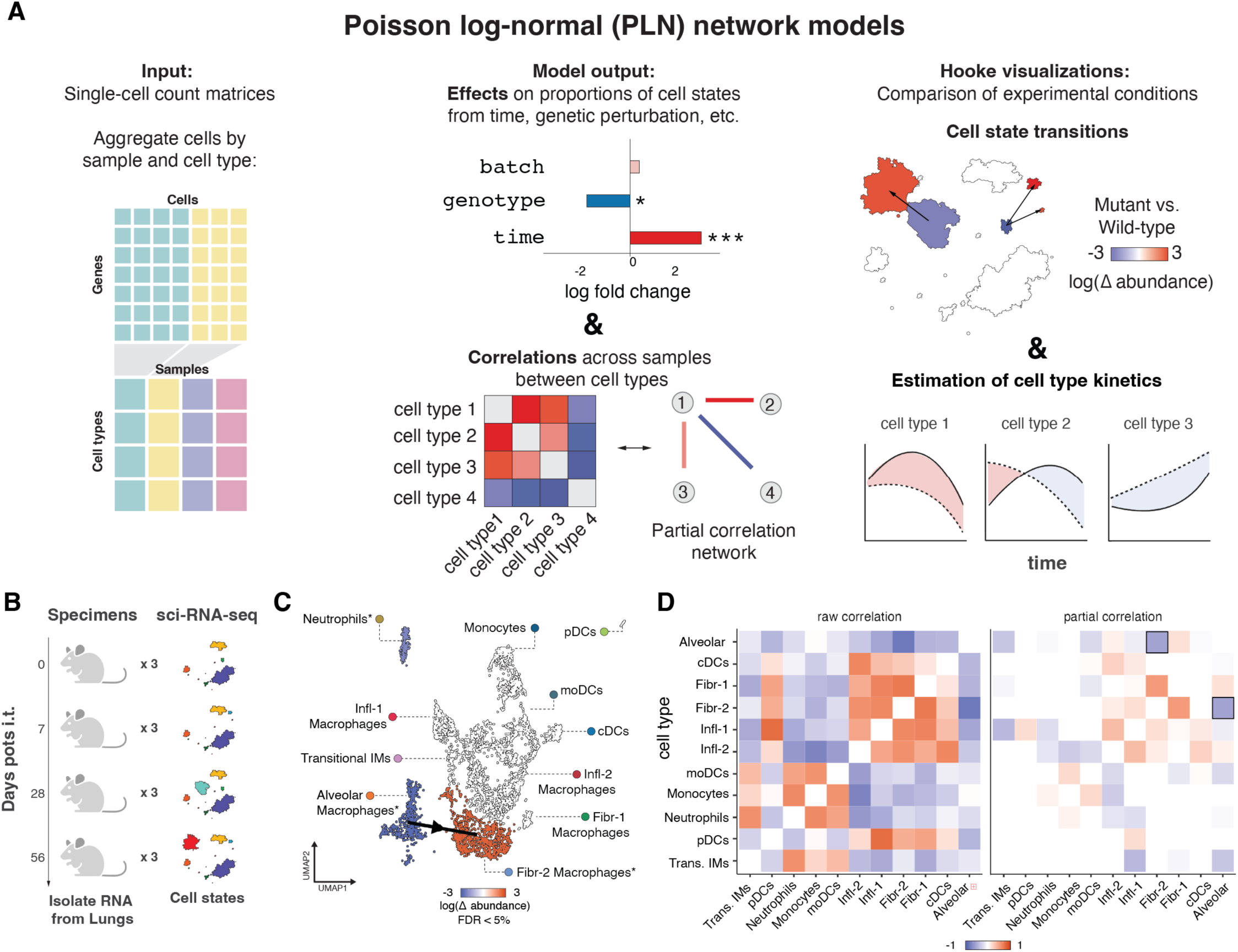
Overview of Hooke. A) Hooke constructs an individual by cell type matrix by tallying the number of cell groups per sample. Poisson-log normal network models estimate the effects on proportions of cell states and partial correlations between cell states. It compares cell state transitions between experimental conditionals and models cell type kinetics. B) A schematic of the experimental design of a mouse model of silicosis. Mice were challenged with intratracheal (i.t.) silica and collected at four timepoints. C) UMAP of myeloid cells colored by fold change of silica exposure relative to wild type (q < 0.05). The arrow indicates a negative partial correlation between a reciprocal fold change between healthy and fibrotic macrophages detected by Hooke. D) Raw correlation between cell states compared to Hooke partial correlations. Boxes surround partial correlations between alveolar macrophages and Fibr-2 macrophages.

Briefly, Hooke works by first estimating the user’s input matrix of cell counts along with metadata for each sample that will be used to model factors in the user’s experiment such as timepoints, drug treatments, genotypes, etc. Then, it estimates per-sample “size factors” that will control for sample-to-sample variation in cell capture levels, similar to methods for library size normalization in RNA-seq^20,22^. Next, it initializes two PLN models: a “main” model that will estimate the proportions of each cell type as a function of all experimental variables relevant to the experiment, along with a second “reduced” model that includes only “nuisance” variances such as batch, etc. Hooke then estimates the coefficients for each model and their confidence intervals using the PLNmodels package. These coefficients reflect the change in the expected counts of each cell type for a given covariate. For the reduced model, Hooke also describes the covariance in cell proportions using variational inference via the PLNmodels package’s optimization routines. These correlations are captured as a network that links pairs of cell types that are still correlated across samples, even after having controlled for changes in all the other cell types (**Figure 1A** and Methods). We hypothesized that the covariation structure of cell type proportions reflects changes over time and alternative fates (i.e. we expect a progenitor and its descendant should be negatively correlated, while cell types that share a common progenitor should be positively correlated).

To illustrate Hooke’s accuracy, we tested its performance by simulating losses of cell types at different effect sizes and embryo sizes and calculating its ability to detect those cell types as differentially abundant (Methods). Hooke performed similar or better than other state of the art methods (**Figure S1A**). We also evaluated Hooke’s ability to detect cell proportion changes between different genotypes as a function of three factors: the abundance of a given cell type, the number of replicates in each genotype group, and the effect size (**Figure S1B**). With 8 specimens per treatment group and 2000 cells per embryo (as in Saunders et al., 2023), large effects (reductions of at least 75%) can be detected for rare cell types such as gill ionocytes and erythrocytes, and modest effects (reductions of ∼10%) could be detected in abundant cell types such as mature fast and slow muscle, showing that Hooke quantifies perturbation effects comparably to state of the art methods for differential composition analysis.

Next, to test our hypothesis that the modeling cell type proportions could link pathological and healthy disease states, we applied it to sci-RNA-seq data from a longitudinal mouse study of silica-induced pulmonary fibrosis^23^, which is a common occupational disease. This data set contains 23,794 nuclei from 12 mouse lungs across four timepoints (0-, 7-, 28-, and 56-days post intratracheal silica) (**Figure 1B**). In the original study, lung macrophages exposed to silica adopted an osteoclast-like transcriptional profile characterized by the expression of bone resorbing proteases and hydrochloric acid. We therefore focused our analysis on the fine-scale cell state annotation of myeloid cells, which contains 11 annotated cell populations (**Figure S1C**).

A Hooke model fit to these data with exposure to silica as a covariate detected a nearly 2.5-fold gain in osteoclast-like, pro-fibrotic macrophages (“Fibr-2” in Hasegawa and Franks et al., 2024), a 3.7-fold reduction in alveolar macrophages, and 2.1-fold reduction in neutrophils (q < 0.05) (**Figure 1C**). The effect sizes reported by Hooke were highly concordant with the Beta-Binomial test used in the original study (**Figure S1D**). Hooke also estimated the correlation structure between cell types in lungs, which was not previously explored. The PLN network linked alveolar macrophages and pro-fibrotic macrophages but not with neutrophils (**Figure 1C,D**). Both macrophage subsets were shown to have the highest enrichment scores of genes associated with osteoclast differentiation, development, and signaling^23^. This link between healthy macrophages and a pro-fibrotic, silicosis-specific state therefore recapitulates the central finding of the original study. Importantly, the link between healthy and osteoclast-like cells was not among the many correlations that the network model was able to attribute to changes in batch or based on changes in other cell types (**Figure 1D**). This analysis demonstrates that, as we hypothesized, the correlation structure between cell proportions captured by Hooke can link disease-specific cell states with their likely healthy counterparts.

### A refined and expanded zebrafish single-cell atlas of perturbed embryos

With a robust method to analyze changes in cell type proportions, we wanted to next apply this approach to investigate developmental processes at scale. To test if Hooke could be applied to atlas-scale data containing multiple timepoints across perturbations, we sought to model cell type abundances during zebrafish organogenesis. In previous and ongoing work, we subjected thousands of embryos to diverse genetic, chemical, and environmental perturbations, followed by whole-embryo single-cell sequencing^20,24,25^. The Zebrafish Single-Cell Atlas of Perturbed Embryos, or “ZSCAPE”, included hundreds of unperturbed or control-injected embryos to use as basis for comparison for phenotyping the perturbations (**Figure 2A**), comprising 1.2 million cells from 18 hours post-fertilization (hpf) to 96 hpf. Each perturbation experiment was projected into this wild type developmental atlas and labels are transferred from their top nearest neighbors in the reference (Methods). This consistent reference space allows us to compare each perturbation’s pattern of differential cell type abundance across the different tissues and cell types in the embryo.

**Figure 2.**
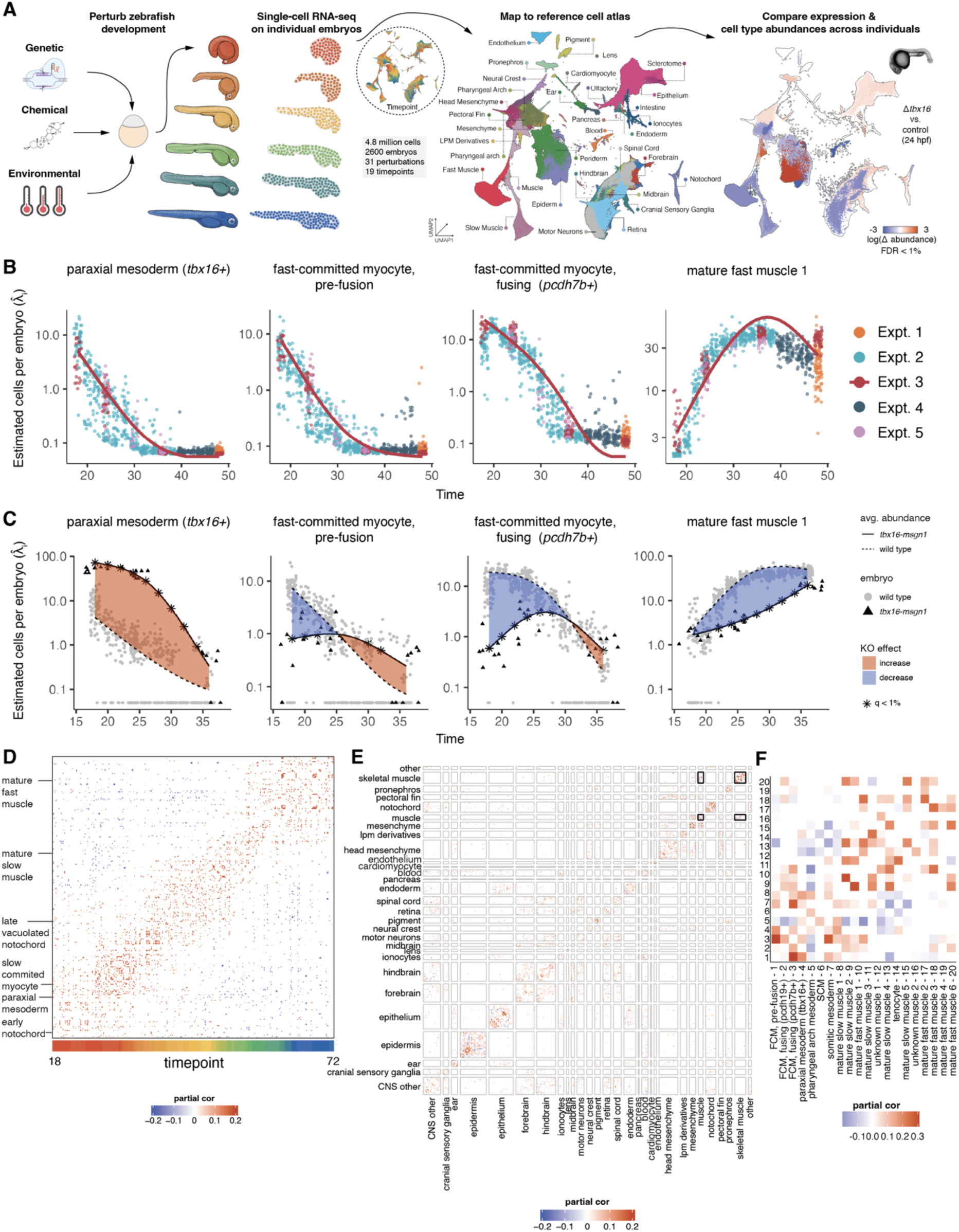
Hooke applied to atlas scale perturbation experiments. A) Schematic of the overall analysis workflow. B) Kinetic curves of wild type fast muscle cell types across multiple experimental batches. The curve plotted is for Expt 3. Points are conditionally predicted per-model embryo estimates (λ_i_) colored by experiment. C) Kinetic plots comparing control to *tbx16-msgn1* in a subset of muscle cell types. The dashed-curve is wild type and the solid curve is *tbx16-msgn1.* Points are conditionally predicted per-model embryo estimates (λ_i_) colored by wild type or *tbx16-msgn1*. * Indicates significance q < 0.01 for a given timepoint comparison between control and *tbx16-msgn1*. D) Partial correlations between wild-type cell types sorted by time of peak abundance. E) Partial correlations between control and *tbx16-msgn1* cell types after accounting for timepoint. The axis is sorted by time within time at peak wild type abundance within projection group. F) Zoom in of muscle and skeletal muscle *tbx16-msgn1* partial correlations. FCM, fast-committed myocyte; SCM, slow-committed myocyte.

Prior to performing an embryo scale analysis of mutant embryos with Hooke, we revised and enhanced our wild type reference atlas. This involved stricter filtering of cells to enhance trajectory resolution and annotation, as well as incorporating approximately 133,000 additional control cells from Barkan *et al*. (Methods), which is referred to here as “reference v2.0”. The UMAP was divided into 30 partitions (“assembly groups”) representing 38 major tissues (**Figure S2A,B)**. The refined UMAP embeddings facilitated the resolution of previously unidentified cell populations, including, among others, specific cell types in the pectoral fin, head mesenchyme, and pronephros.

Despite these improvements, resolving cell type annotations in the central nervous system (CNS), particularly among progenitor states, remained challenging. To address this, we applied the Tangram algorithm to align our single-cell data with a Stereo-seq dataset from 24 hpf zebrafish embryos^26,27^. This approach can help infer the spatial origin of cell types with similar transcriptomes that would otherwise be difficult to disentangle^2^. This alignment enabled the separation of CNS progenitors into their respective forebrain, midbrain, hindbrain, and spinal cord assembly groups **(Figure S2D-E**). Within each assembly group, cells were further sub-clustered and annotated based on the expression of known marker genes. The earlier atlas included 156 fine-level cell type annotations (“cell type sub”), 95 broadly labeled cell types (“cell type broad”), and 37 tissue types (**Figure S2C**). Following recombination and re-embedding of these assembly groups, and including our new sequencing data, we identified 163 additional cell types at the finest resolution in reference v2.0 (**Figure S2E**), resulting in a total of 319 distinct cell types and 174 broad cell type categories (**Table S1**).

### An embryo-scale Hooke model of cell abundances captures their kinetics and shared cell abundance phenotypes

Examining the effects of gene knockouts at a single timepoint provides a limited understanding of gene function, as the importance of a gene may vary across developmental stages, and tissues may respond differently over time. We first sought to verify that Hooke can model the kinetics of cells as they develop in the embryo, even in single-cell datasets comprised of many timepoints and multiple batches. Fortunately, the PLN framework can include covariate terms, such as experimental batch or sampled timepoint, when modeling changes in cell abundance.

The ZSCAPE dataset contains 19 timepoints and five experimental batches, so we fit a model that included timepoint and experimental batch as covariate terms. This model was able to describe the smooth kinetics of cells in the embryonic skeletal muscle as they transition through different stages from *tbx16+* paraxial mesoderm, through myoblasts and fusing myocytes, to mature muscle (**Figure 2B**). To test the model’s ability to subtract batch effects, we predicted each cell type’s relative abundance in each embryo conditional on its observed counts. After using the model to account for batch effects, the estimated proportions for each cell type were in agreement, even embryos from different batches and experiments, and they followed the overall kinetic trend of the model (**Figure 2B**). Importantly, the curves spanned the full extent of our atlas, from 18 hpf to 96 hpf, despite the fact that the five experiments covered distinct time intervals, sampling regimes, and levels of cell capture and sequencing depth (**Figure 2B**). This demonstrates that Hooke can capture the times when different cell types emerge or reach their peak abundance, even when integrating data from different experiments or batches.

We next sought to use Hooke to study individual genetic perturbations as a means of defining the genetic requirements for cell types throughout the embryo. The ZSCAPE data contains 23 zebrafish G0 knockouts generated by CRISPR-Cas9 mutagenesis (crispants). Because each perturbation in ZSCAPE was collected at multiple timepoints, we can also assess the kinetics of perturbed cell types by fitting a Hooke model with genotype and timepoint as covariates (**Figure 2C**). By comparing the two kinetic curves, we can detect the time interval within which a specific cell type begins to be lost or stalled in a crispant. We performed this analysis in crispants lacking both *tbx16* and *mgsn1*. These transcription factors regulate the differentiation of the mesodermal lineage from the neuromesodermal progenitor cells (NMPs) that normally give rise to two cell lineages: somitic muscle and spinal cord neurons^28^. Hooke’s model detected that embryos lacking *tbx16* and *msgn1* had reduced mature muscle and a significant accumulation of cells in the earlier progenitor state (**Figure 2C**), consistent with our expectation that mesodermal progenitors will fail to mature in this crispant^29^.

Having verified that Hooke can locate the cell types and times within which mutations exert their effects, we scaled this approach and modeled genotype and time across our complete perturbation set of 23 crispants^20^. We found that 308 of the 319 cell types were significantly differentially abundant cell types (DACTs) in at least one perturbation and timepoint (**Table S2**). Previous modeling strategies limited us to comparing cell type gains and losses at a single timepoint, as the Beta-Binomial allows for modeling a single outcome at a time.

However, because the PLN framework allows for simultaneous modeling of how different predictors influence different count outcomes, we summarized DACTs across the atlas time range, allowing us to visualize the cell-type specific effects of each perturbation across the whole time series at once (**Figure S1E**). Hierarchical clustering of these fold changes resulted in extensive phenocopying amongst crispants for mesoderm lineage factors (*cdx4*, *cdx1a*, *tbxta*, *tbx16*, *tbx16l*, *msgn1*, *wnt3a*, *wnt8a*, *noto*, *smo*), neural crest lineages (*foxd3*, *tfap2a*), and two clusters of central nervous system lineage factors (1: *egr2b*, *epha4a*, *hoxb1a*, *mafba*, *zc4h2*; and 2: *phox2a*, *foxi1*, *hgfa*, *met*).

We next tested the hypothesis that correlations between cell types’ proportions during embryonic development reflect their underlying lineage relationships. Running Hooke on 107 wild-type zebrafish embryos from 18 to 72 hpf showed that cell types tend to be positively correlated with cell types present in the same time window, reflecting their synchronous development, and negatively correlated with cell types present at much later time windows (**Figure 2D**). The latter observation is consistent with early progenitors ultimately giving rise to differentiated tissues. However, early cell types might be anti-correlated with ones that develop from unrelated lineages, so we next examined links within and across tissues in a Hooke model fit on embryos lacking *tbx16* and *msgn1*. We fit the model to both *tbx16-msgn1* crispants and paired non-targeting controls in a manner designed to subtract the changes over time shared by both genotypes but capture any differences between genotypes in the correlation structure. This model revealed strong links within tissues and much weaker correlation structure between different tissues **(Figure 2E**). Moreover, links between co-varying cell types were especially pronounced in tissues involved in the *tbx16-msgn1* phenotype and often reflected lineage relationships. For example, paraxial mesoderm (*tbx16+*) cells were inversely proportional to mature fast muscle **(Figure 2F**). In contrast, states derived from the same progenitor (e.g. distinct subsets of mature slow muscle) tended to be positively correlated **(Figure 2F**). These analyses support our hypothesis that the correlation structure in cell proportions of the developing embryo reflects lineage relationships between them. Furthermore, the patterns of partial correlation structure between cell types can be used to infer how cell types co-vary in their abundances over time.

### Differential analysis of mutant embryos defines the genetic requirements of embryonic cell types

In constructing ZSCAPE, our goal was to define how cell types depend on one another and on key genes in the genome for their development. To illustrate Hooke’s ability to reveal genetic requirements of cell types, we applied it to study the zebrafish pronephros. The pronephros, or the embryonic kidney, is composed of two nephrons running in parallel along the anterior-posterior axis of the embryo and is responsible for the removal of metabolic waste^30^. At the 28 somite (23 hpf) stage, it is segmented into nine distinct morphological and functional parts, each responsible for a specific function. In our reference and perturbation datasets, we captured 32,840 pronephros cells (approximately 15 cells/embryo). Canonical gene markers were used to annotate the podocytes (*wt1a, wt1b, nphs1, nphs2*), multiciliated cells (*odf3b, jag2b, dzip11*), proximal straight tubule (*trpm7, slc13a1*), early distal tubule (*slc12a1*), late distal tubule (*clcnk, mecom, slc12a3, pppr1b*), proximal convoluted (*slc4a4a, slc4a2a, slc26a2, slc20a1a, pdzk1, slc20a1a*) and corpuscles of Stannius (*stc1l*). In addition to these 7 segments, we also identified a population of renal progenitor cell types (**Figure 3A**).

**Figure 3.**
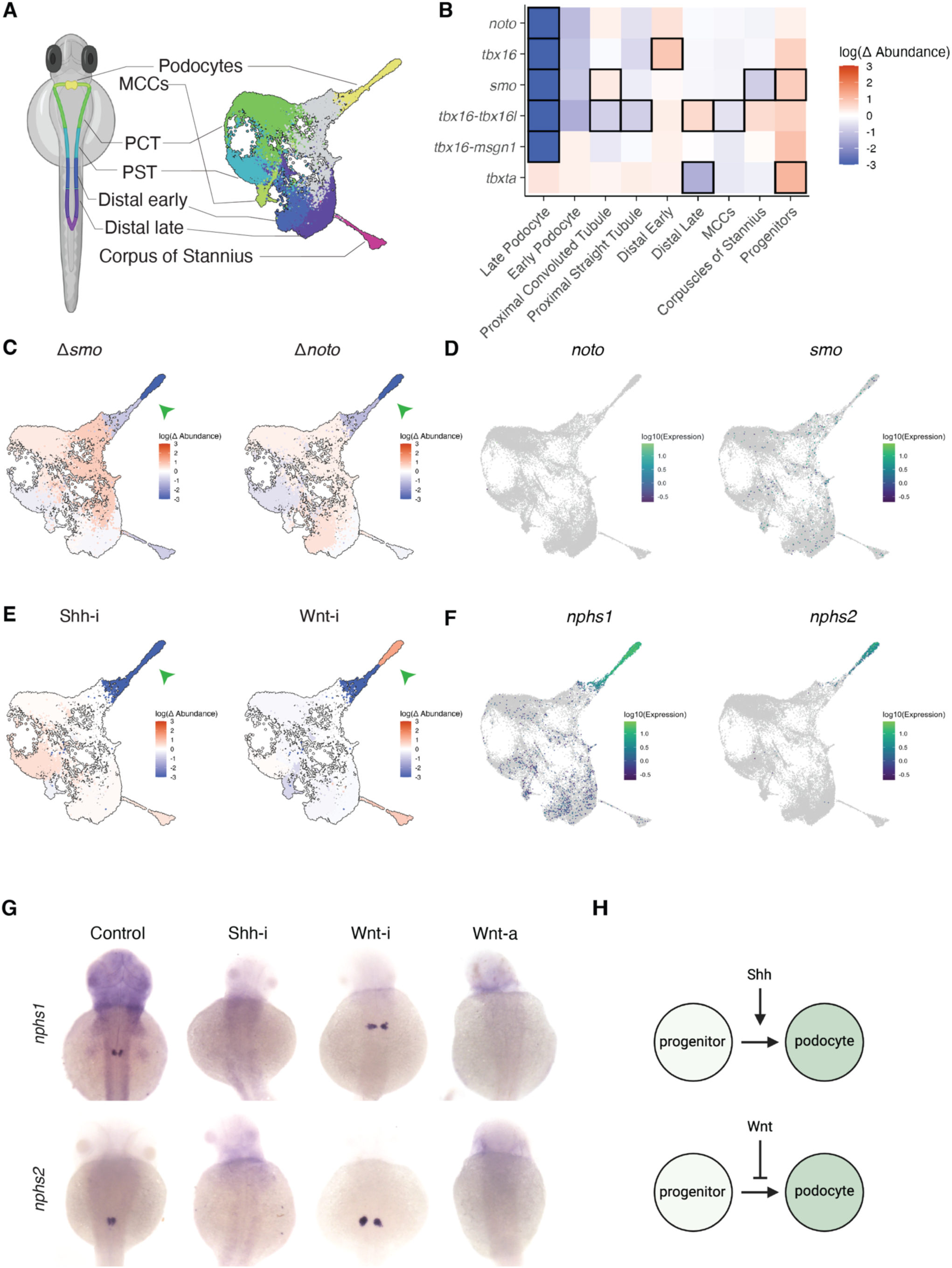
Regulation of podocyte development by Shh and Wnt signaling. A) UMAP showing developing kidney cell types along with their arrangement in the pronephros. Renal progenitor cells are colored gray. B) Hierarchical clustered heatmap of Hooke reported cell type abundance changes in pronephros clusters and their corresponding cell types at 36 hpf. Boxes indicate significant changes (q < 0.05). LP, late podocyte; EP, early podocyte, MCC, multi-ciliated cells; PCT, proximal convoluted tubule; PST, proximal straight tubule; DL, distal late; DE, distal early; CS, corpus of stannius. C) UMAP colored by changes in each cell type’s abundance in smo and noto crispants at 36 hpf as estimated by Hooke. D) Expression of markers in the Kidney UMAP colored by log10 of gene expression of *noto* and *smo.* E) UMAP colored by changes in each cell type’s abundance in Shh-inhibited (Shh-i) and Wnt-inhibited (Wnt-i) embryos at 36 hpf. Shh-i added at 6 hpf (shield stage) and Wnt-i added at 13 hpf (8-somites stage). F) Expression of markers in the Kidney UMAP colored by log10 of gene expression of *nphs1* and *nhps2*. G) WISH of control, Shh inhibited, Wnt inhibited, and Wnt-activated (Wnt-a) treated embryos stained for podocyte-specific genes (*nphs1*, *nphs2*) at 48 hpf. Shh-i added at 6 hpf (shield stage) and Wnt-i added at 13 hpf (8-somites stage). H) Diagram depicting our proposed model of signaling pathway regulation of the podocytes.

How genes and signals control the development of each of these important segments is still not fully understood. We sought to explore the requirements of pronephric cell types using Hooke. To test whether subsets of pronephric cells require any of the genes we previously perturbed, we fit a Hooke model across all the crispants using genotype and time as covariates in the model. This analysis revealed significant changes in distinct pronephric cells of several crispants, including *smo*, *noto,* and *tbx16* (**Figure 3B,C**).

One key feature we observed across genotypes was the differential loss of late podocytes, which play an important role in the glomerular blood filtration barrier^31^. A deficiency in *tbx16* has been previously associated with a reduction in podocytes and proximal tubule segments^32^. The loss of the paraxial mesoderm in *tbx16* mutants has been shown to disrupt the expression of a retinoic acid (RA) synthesis gene, suggesting that paraxial mesoderm is a key source of RA for nephron patterning^33^. However, other signaling pathways may be involved in the effects observed in the *smo* and *noto* crispants, which have not been previously described to have a depletion of podocytes. *Smo* is the receptor for Sonic hedgehog (Shh) signaling whereas zebrafish *noto* mutants and crispants lack a notochord^34,20^. Importantly, the notochord serves as a midline signaling source of Shh that directs the patterning of surrounding tissues^35^. Since *noto* and *smo* genes are not expressed in the pronephric cell types (**Figure 3D**), this suggests a cell-nonautonomous effect of these genes on the pronephros^34,36^.

We hypothesized that Shh signaling plays an important role in podocyte development, as the pronephros is also located in the midline of the zebrafish embryo, adjacent to the notochord. To test this hypothesis, we examined the data from Barkan *et al.* in which we applied sci-Plex to hundreds of embryos exposed to signaling pathway inhibitors during organogenesis (BMP, FGF, Notch, RA, Shh, TGFβ, and Wnt)^25^. These embryos were collected at overlapping timepoints with those examined in Saunders *et al.* (2023), with 8 embryos per timepoint and matched controls. We fit a Hooke model to the Barkan *et al*. dataset across the entire embryo using chemical inhibitor and timepoint as covariates. For this analysis, we specifically focused on the differential abundance patterns in the pronephros. We compared the chemically inhibited effect sizes to those with genetic perturbations at 36 hpf. Shh-inhibited embryos (cyclopamine) had similar abundance changes to the *smo* and *noto* embryos, including a 97% reduction in podocytes (**Figure 3C,E**). Conversely, in Wnt-inhibited (Wnt-59) embryos, the podocytes had a 4-fold increase in abundance, suggesting Wnt is a negative regulator of podocyte development. We performed a series of whole-mount in situ hybridization (WISH) experiments to validate these changes in podocyte abundance using the previously identified markers *nphs1* and *nphs2*^37^, and observed a lack of podocytes when Shh was inhibited and an increase in podocyte expression when Wnt was inhibited (**Figure 3F,G**). We also tested a Wnt-activator (BIO) and saw an absence of podocytes supporting the idea that Wnt is a negative regulator of podocyte development (**Figure 3G**). Based on these concordant results, we propose a model of Shh positive regulation and Wnt negative regulation of the podocytes (**Figure 3H**).

### Covariation across individuals resolves molecular transitions in the developing embryo

Although our initial analysis of ZSCAPE detected many cell types that were lost in each crispant^20^, the lack of a high-resolution map of the lineage relationships between these cell types precluded us from clearly delineating which cell types directly required each perturbed gene. Single-cell “trajectory analysis”, in which transcriptionally similar cells are assumed or inferred to be developmentally related, is often used to study how cells regulate and are regulated by genes as they develop. However, such inference can mistakenly link cell types that are transcriptionally similar but not lineally related^38^. We reasoned that Hooke’s models of how cell types co-vary with one another across time and in response to perturbations could be used to generate accurate maps of lineage relationships between cell types that could be used to better interpret perturbations.

To explore this idea and better understand how cell types arise from other cell types during embryogenesis, and how they might share genetic requirements, we next developed “Platt”, an algorithm that uses Hooke’s cell abundance models to infer cell state transitions even in very complex tissues or hard to resolve single-cell trajectories (**Figure 4A** and Methods). Platt is named for Julia Platt, who first suggested the neural crest origins of the skull ^39^. Platt first subclusters cells into “microstates” and fits a Hooke model on wild type cells over time, estimating changes in cell proportions and recovering a partial correlation network among microstates. Next, Platt initializes a “pathfinding graph” over which cells could plausibly travel as they transition between microstates. The pathfinding graph is constructed by linking microstates that are both transcriptionally similar and deemed partially correlated by Hooke. Then, Platt orients this graph so that cells travel over the shortest paths from microstates common in early embryos towards microstates common in later embryos. Next, Platt considers any available perturbation experiments, reasoning that a perturbation that depletes cells in an “ancestral” microstate should also deplete cells in any “descendant” microstates. For each putative transition, Platt checks whether the two states were concordantly gained or lost in at least one genetic perturbation, assigning weights and labels to each edge based on this support. Finally, the graph is pruned, eliminating any weakly-supported edges needed to ensure that it is acyclic.

**Figure 4.**
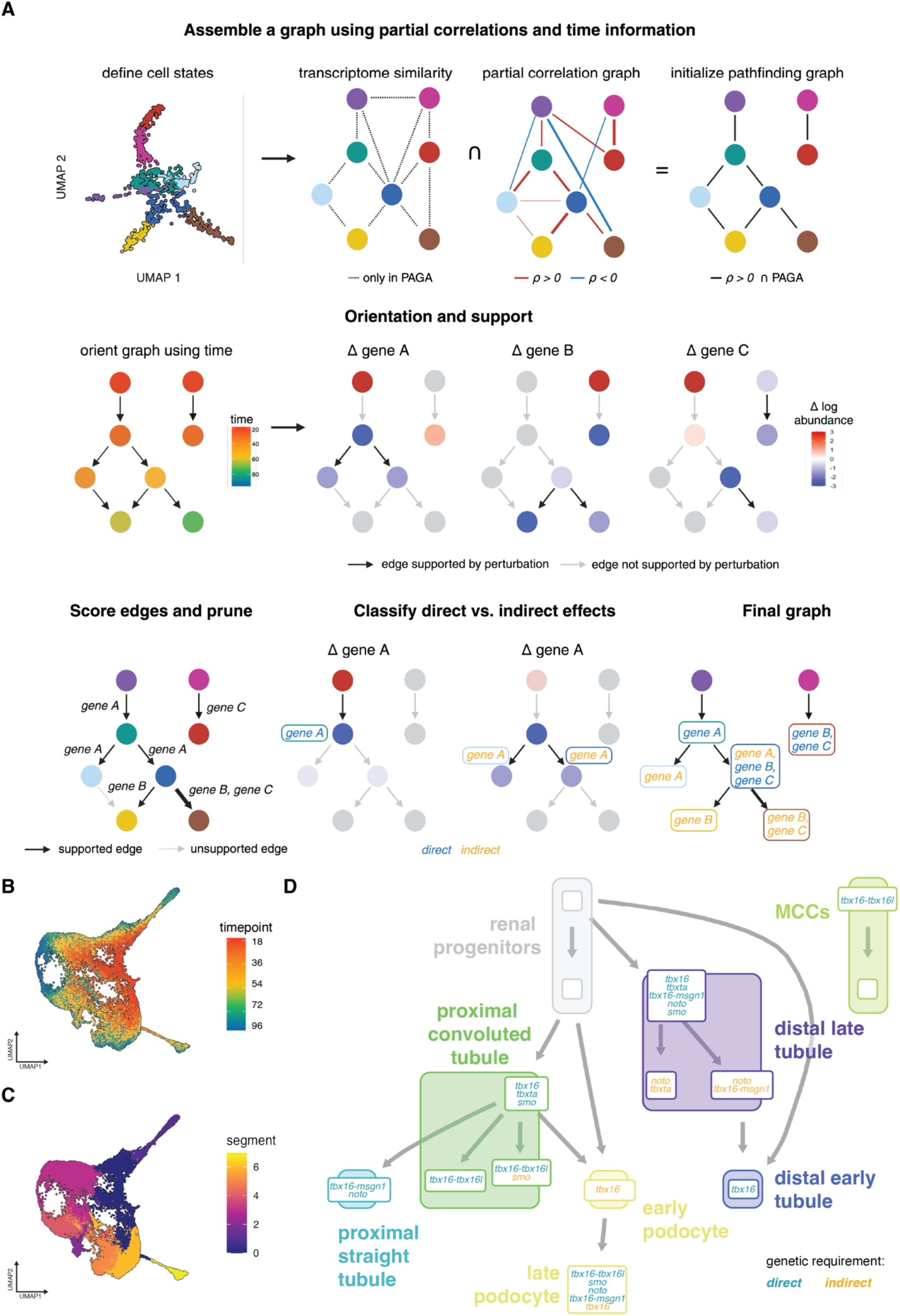
Platt builds cell state transition graphs using time series and perturbation data. A) A cartoon UMAP and schematic of the Platt cell state transition graph assembler algorithm. PAGA: Partition-based graph abstraction. B) Sub-UMAP of the kidney colored by timepoint. C) Sub-UMAP of the kidney colored by spatial segment. D) A Platt-derived cell state transition graph built using wild type and mutant data. The smaller boxes denote clusters. Larger colored boxes denote cell types and match colors in Figure 3A. Cluster nodes are labeled by genetic requirements (subset to the perturbations in Figure 3) and colored by their direct (blue) or indirect (yellow) classification.

As a proof of principle, we applied Platt to pronephros cells from wild-type and genetically perturbed zebrafish embryos in ZSCAPE. Despite their specialized functional roles, pronephric cells express many genes in common, and accordingly do not develop along a simple linear or branched pseudotime trajectory (**Figure 4B,C**). The resulting state graph linked microstates present within each spatial segment of the pronephros of early embryos to microstates present in the corresponding spatial segment of later embryos (**Figure 4D**).

Moreover, Platt judged the links within the segments as supported by perturbations, whereas it reported links between segments as lacking direct perturbational evidence. For example, perturbations Platt assessed to support links between microstates included *tbx16-msgn1* and *tbx16*-*tbx16l.* Tbx16 is likely required by the pronephros via retinoic acid signaling^32^. Taken together, these analyses demonstrate that Hooke and Platt can work together to resolve complex cellular trajectories on the basis of shared genetic requirements in embryonic tissues.

### A whole-embryo map of molecular state transitions during zebrafish organogenesis

Having established algorithms for inferring lineage relationships between cell types based on their shared genetic requirements, we next applied Platt to ZSCAPE to construct an embryo-scale map that spanned organogenesis in the zebrafish (**Figure 5A**). Platt linked 153 cell types (48% in the atlas) into the resulting state graph, with 173 transitions in total. 110 of the cell types in the reference atlas had at least one direct “ancestral cell type” in the graph. Cell types that emerge prior to 18 hpf (the earliest timepoint represented in ZSCAPE) were frequently found at component roots, and cell types that emerge later found at the leaves (**Figure S3A**). Cell types with neither ancestors nor descendants tended to be much lower in proportion and were commonly annotated as potential doublets based on markers (**Figure S3B**). However, Platt did successfully incorporate very rare or specialized cell types, such as pectoral fin cleithrum (∼2-3 cells recovered per embryo).

**Figure 5.**
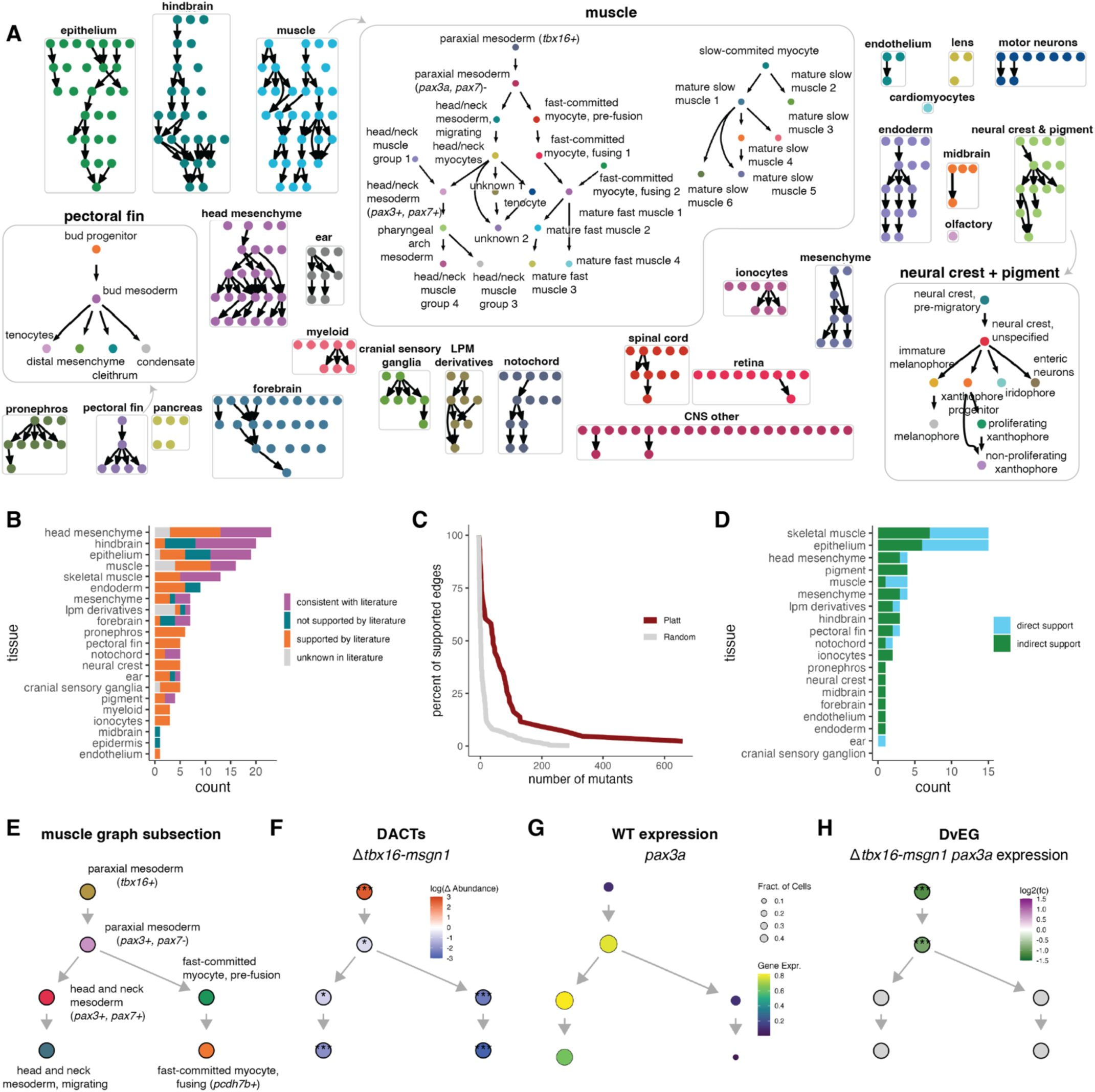
A whole-embryo map of molecular state transitions. A) State transition graphs for each assembly group with zoom-in graphs for pectoral fin, muscle, and neural crest and pigment. B) The count of literature support in each tissue graph. “Consistent with literature” indicates the Platt graph cell type resolution is finer than what is reported. C) The cumulative sum of edges supported by Monarch genetic requirements in the Platt graph compared to 100 randomized graphs. D) The count of direct or indirect support from perturbations in each tissue graph from Saunders *et al*., 2023. E) A top subsection of the muscle graph labeled by cell type. F) The muscle Platt graph colored by log cell abundance fold change (*tbx16-msgn1*/control) as estimated by Hooke (*: p < 0.05, **: p < 0.01, ***: p < 0.001). G) Platt graph colored by *pax3a* wild type expression. H) Platt graph colored by log fold change (*tbx16-msgn1*/control) of *pax3a* in the *tbx16-msgn1* mutant (*: p < 0.05, **: p < 0.01, ***: p < 0.001).

Platt’s map of cell state transitions between zebrafish embryonic cell types was rich with support from literature; 84.3% of links were observed or inferred in previous studies (**Figure 5B** and **Table S3**). However, because cell type definitions and boundaries are not consistent across past studies, we sought to systematically assess the quality of the graph. First, we manually matched each ZSCAPE cell to corresponding types in the ZFIN Cell Ontology, which enabled us to look up all genes known to be required for that cell type according to the Monarch Initiative^40^ (Methods). Next, we asked whether two cell types linked by Platt both shared a common genetic requirement from Monarch. Of 173 transitions in Platt’s graph, 87% were supported by at least one shared genetic requirement, with many edges supported by many genes (**Figure S3C,D**). We compared the Monarch support of our Platt graph edges to 100 randomly generated graphs (**Figure 5C** and **Figure S3C**). Thus, while the Platt state graph is not fully connected, the edges that are present are well-supported by prior studies and highly consistent with the consensus view of how cell types develop from one another in the zebrafish embryo.

We next explored how our own perturbations supported each transition as a means of understanding how our choices of target genes illuminated transitions in different tissues. Scoring each edge for support from each ZSCAPE perturbation revealed 23 edges (14%) were supported by at least two independent perturbations.

Many supported edges, but not all, were concentrated in skeletal muscle, the central nervous system (CNS), and the neural crest derivatives, consistent with the fact that Saunders *et al.*, 2023 targeted many transcription factors important in these lineages (**Figure 5D**). A further 32 edges (19%) were supported by just one perturbation, while 40 edges (24%) were supported indirectly, leaving the remaining 70 edges (42%) consistent with time but otherwise without support from perturbations on each side of the corresponding state transitions (**Figure S3E**). Upstream of these direct cell losses, 12% have a significantly enriched ancestor, likely corresponding to a block in that cell type’s further development.

We next examined the genes that change as cells transition over the graph by performing systematic differential gene expression analysis over the map. We iterated over the map and, for each node, looked for several patterns of differential expression (**Figure S4A**). For example, “terminal selector” and “multilineage priming (MLP)” genes might activate specific fates, genes expressed in progenitors only might be required for maintaining the progenitor states, and genes excluded from certain fates might repress that fate^38,41–43^. Of the 2350 classed as transcription factors by CIS-BP^44^, 1739 were expressed in a manner suggestive of such fate-determining roles in one or more of 92 cell types. Terminal selectors included *mitfa*, *phox2a*, and *wt1a* in melanophores, epibranchial ganglion, and podocyte cells, respectively (**Figure S4B-D**). The MLP effect of *pax3a* was observed in paraxial mesoderm (*pax3a, pax7-*) progenitors as they commit to either head and neck mesoderm or fast muscle fates, consistent with its known role in governing these decisions^45^ (**Figure S4E**). We next asked which of the genes were known to be important for the cell types in which we detected their regulation. Of the 1739 transcription factors, 32% were directly required by at least one cell type as reported in the Monarch Initiative Knowledge Graph^40^, ZFIN, or Gene Ontology Biological Process databases (Methods). Furthermore, 40% of cell types had a significant overlap between Platt-determined requirements and Monarch-determined requirements (**Figure S4F,G)**. These requirements were broadly distributed over the Platt state graph, demonstrating that dynamically expressed transcription factors are often required by the cell types that regulate them.

We reasoned that perturbations that disrupt the differentiation of some cell types might also disrupt gene regulation in their progenitor cell types, so we next asked whether cell type losses were “anticipated” by transcriptional phenotypes in the upstream transcriptional states. We performed a differential expression analysis comparing cells from perturbed embryos to the same cell type from control embryos (Methods). Across all 24 genotypes, we uncovered 21,119 differentially expressed genes (DEGs). However, DEGs were detected in many cell types that were not significant DACTs in the corresponding perturbation (**Figure S4H**). In fact, few DEGs were detected in cell types substantially lost in a perturbation, likely reflecting that, without adequate cell sampling in both conditions, power for transcriptional comparisons is limited. However, DEGs were commonly detected in cell states upstream of DACTs. We then asked which genes are “deviantly expressed” (DvEG) in each perturbation, defined as genes that are under- or overexpressed in perturbed cells undergoing a transition relative to controls (**Figure S4I**). For example, *tbx16*-*msgn1* crispants fail to generate skeletal muscle properly and *pax3a*, a key regulator of myogenesis, was deviantly expressed in myogenic progenitors (**Figure 5E-H**). In total, we detected 12,099 DvEGs across the ZSCAPE perturbation collection, including in cell types present at normal wild type abundance. Of these DvEGs, 795 (7%) were themselves transcription factors, and 1297 (11%) were known genetic requirements of the cell types in which they were deviantly expressed, and 2,169 genes (18%) had phenotypes affecting the associated tissue. Together, these analyses underscore the utility of an embryo-scale state graph for interpreting the transcriptional and cellular effects of perturbations and its potential for nominating future targets.

### Graph-assisted identification of novel genetic requirements

To expand Platt’s map and recruit unlinked cell types or transitions with scarce support, we next sought to use the map to predict new genetic requirements that could be targeted in perturbations. Amongst the genes Platt associated with many state transitions and fate decisions was *lmx1bb* (**Figure S4J**), which has been previously implicated in the development of the kidney, eye, limb, ear, and midbrain-hindbrain boundary^46,47–48^. We generated G0 knockouts (crispants) targeting *lmx1bb* and its paralog, *lmx1ba*^49^, using CRISPR-Cas9 mutagenesis (*Δlmx1b*, Methods). WISH confirmed the loss of podocytes and the midbrain-boundary, recapitulating previous studies (**Figure S5A,B**). To comprehensively define the molecular phenotype of *Δlmx1b,* we collected eight replicates of *Δlmx1b* and paired injection controls at six timepoints across organogenesis: 18, 24, 36, 48, 60, and 72 hpf (**Figure 6A** and **Figure S5E**). We then used sci-Plex to profile these cells, where embryo-specific barcodes were used to label each sample’s perturbation and timepoint of collection. In total, 572,958 cells were collected from 168 embryos, with an average of ∼3,400 cells captured per embryo and ∼660 UMIs per cell (**Figure S5C,D**). We projected the data onto our reference atlas and transferred cell-type labels using a nearest-neighbor approach (Methods).

**Figure 6.**
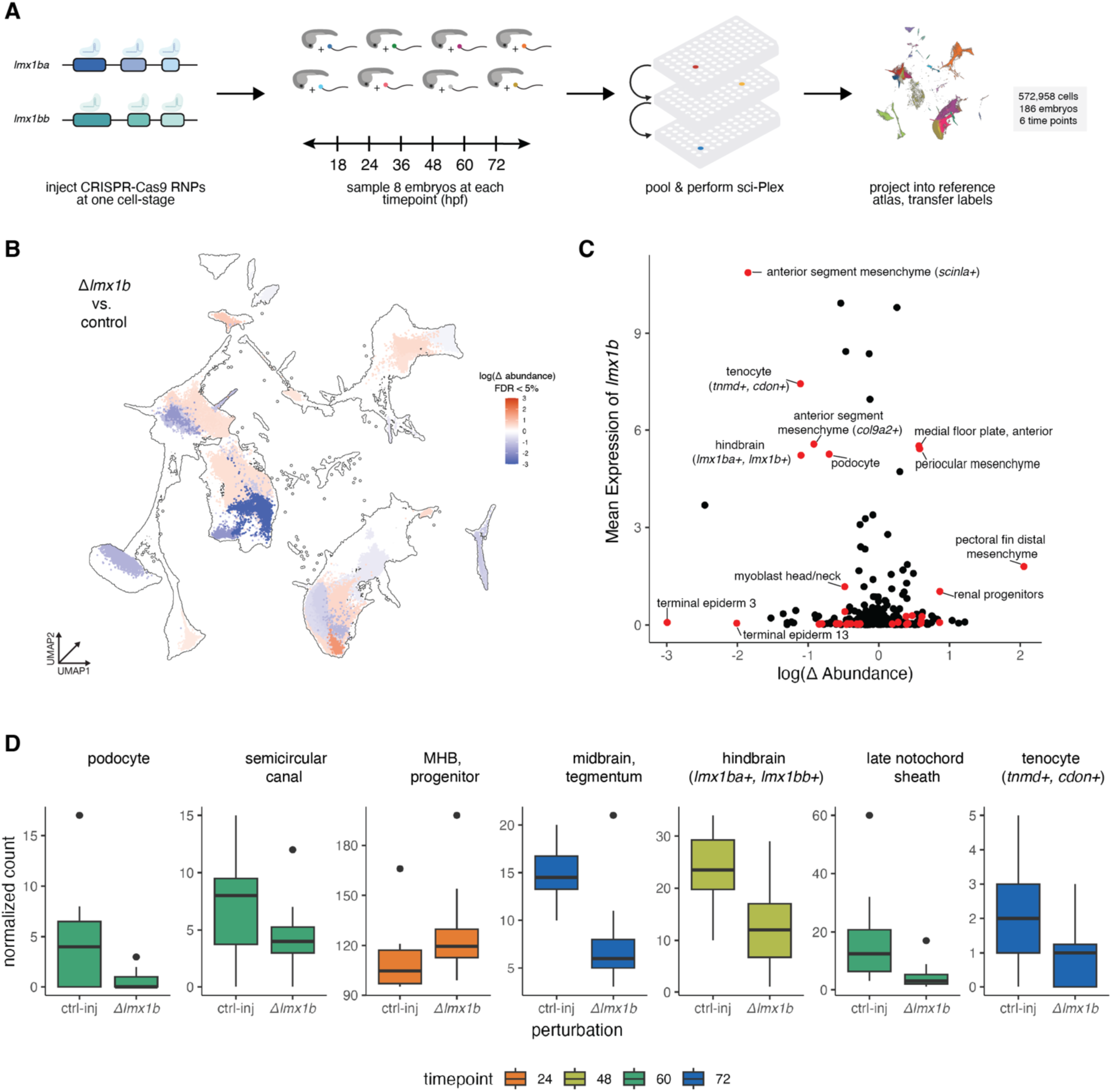
Characterizing the *Δlmx1b* crispant. A) A schematic of the experimental design. B) Global UMAP colored by changes in each cell type’s abundance in the *Δlmx1b* crispants at each cell type’s peak abundance as estimated by Hooke (q < 0.05). C) Scatterplot comparing the mean expression of *lmx1b*+ cells compared to the relative abundance of *Δlmx1b* vs control embryos in each cell type. D) Boxplots of normalized cell counts from wild type vs. *Δlmx1b* crispants from select cell types colored by timepoint. MHB: midbrain-hindbrain boundary.

To phenotype the *Δlmx1b* crispant, we first fit a Hooke model using genotype and time as covariates. Hooke detected 109 significant DACTs across the embryo over the six timepoints (q < 0.05) (**Figure 6B** and **Figure S5F**). Some cell types with detectable expression of *lmx1b* were unaffected or even significantly more abundant in crispants than controls, while other cell types with undetectable *lmx1b* expression were significantly differentially abundant in the crispants (**Figure 6C**). As expected, Hooke detected significant losses in podocytes, the semicircular canal (ear), the midbrain, and the hindbrain (**Figure 6D**). We also observed losses in cell types not previously reported to depend on *lmx1b*, such as the late notochord sheath and an unknown tenocyte population (**Figure 6D**). We reasoned that the indirect effects of disrupting *lmx1b*, in which descendants of cell types that express and require the genes were also lost, might be frustrating our efforts to locate the cell types that directly require *lmx1b*.

### Platt-guided differential expression analysis locates putative genetic requirements

To deconvolve indirect and direct effects of disrupting *lmx1b* in the zebrafish embryo, we used Platt to scrutinize the cell abundance phenotype of *lmx1b* over our embryo-scale map of cell state transitions. We focused on DACTs in lineages where *lmx1b* was identified as a selectively activated gene and resulted in a significant abundance change (**Figure 7A-C**). Painting each graph by abundance changes at the timepoint of each cell type’s peak abundance in the wild type pinpointed where losses begin and how they propagate through subsequent transitions (**Figure 7B**). Notably, cell types that specifically expressed *lmx1b* in the wild type were lost in the kidney (podocytes), notochord (vacuolated cells), and head mesenchyme (tenocytes (*tnmd+, cdon+*)), anterior segment mesenchyme (*col9a2*+) and anterior segment mesenchyme (*scin1a*+)) (**Figure 7B,C**). The notochord sheath cells were also lost, even though they did not normally express *lmx1b* at high levels (**Figure 7B**), potentially reflecting a non-autonomous requirement for *lmx1b*. Moreover, ancestors of these depleted cell types such as the renal progenitors and periocular mesenchyme were significantly enriched, consistent with their failure to progress into the podocyte or anterior segment mesenchyme fates, respectively. Visualizing and interpreting the differential abundance analysis of the *Δlmx1b* crispants in the context of the Platt state graphs thus provided a potential explanation for why some *lmx1b*-expressing cell types were lost as others were gained. In total, analyzing *Δlmx1b* crispants lent experimental evidence to four state transitions, two of which were previously unsupported by any previous ZSCAPE perturbation (**Figure S6**), demonstrating that Platt can locate novel genetic requirements and use them to reinforce its own maps of how cell type depend on one another.

**Figure 7.**
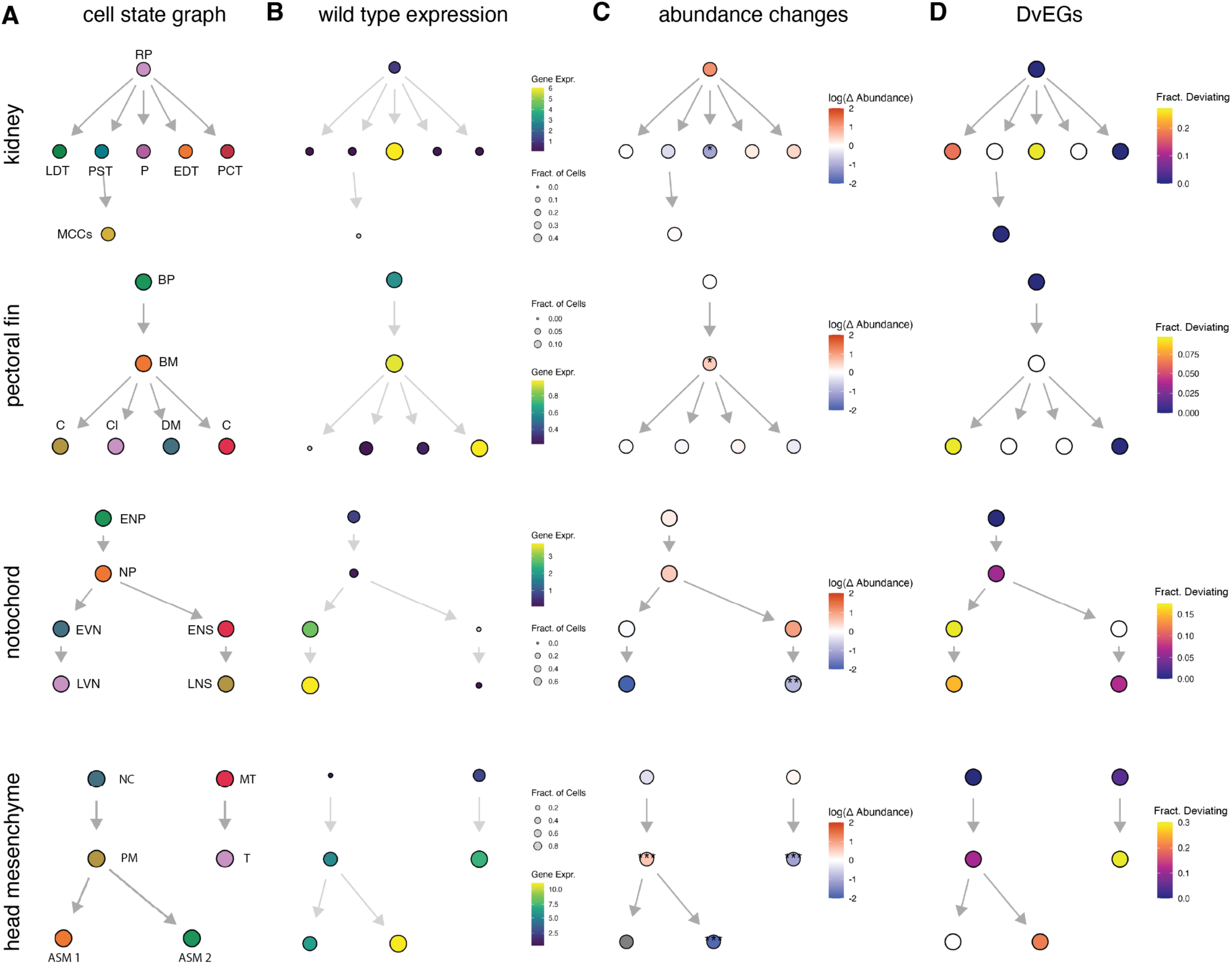
Platt graphs help organize changes in abundance and gene expression. A) Platt-derived state graphs annotated by cell type. Colors represent cell types: RP, renal progenitors; PST, proximal straight tubule; P, podocyte; PCT, proximal convoluted tubule; EDT, early distal tubule; LDT, late distal tubule; MCCs, multiciliated cells; BP, bud progenitor; BM, bud mesoderm; C, condensate; DM, distal mesenchyme; Cl, cleithrum; T-tenocyte; ENP, early notochord progenitor; NP, notochord progenitor; EVN, early vacuolated notochord; LVN, late vacuolated notochord; ENS, early notochord sheath; LNS, late notochord sheath; NC, neural crest, periocular mesenchyme-fated; PM, periocular mesenchyme; ASM 1, anterior segment mesenchyme (*col9a2*+); ASM 2, anterior segment mesenchyme (*scinla+*); MT, mesenchyme, tenocyte-like (*tnmd+*); T, unknown (tenocyte, *tnmd+, cdon+*) B) Platt-derived state graphs colored by expression of *lmx1b*. C. Platt-derived state graphs are colored by significant log fold changes in the abundance of each cell type (*: p < 0.05, **: p < 0.01, ***: p < 0.001). D) Platt-derived state graphs colored by the fraction of significant (q < 0.05) DvEGs in each cell type.

To further investigate the mechanisms underlying cell types that require *lmx1b*, we explored the transcriptional responses of each cell type to the *Δlmx1b* crispant. Pairwise testing between pseudo-bulked control and *Δlmx1b* crispant cells revealed 1,748 unique DEGs across all cell types and 12,099 DvEGs across all cell types in the embryo. These included numerous components of the extracellular matrix (ECM), consistent with a previous study in mice that reported ECM components as direct targets of *lmx1b*^50^. For example, DvEGs included *col2a1a* (**Figure S7A,B**), which was underexpressed along the transitions from notochord progenitor cells to sheath cells and from otic support cells to the semicircular canal, where *lmx1b* is known to promote the expression of other ECM components^46^. Two teneurin genes, highly conserved transmembrane glycoproteins^51^ and which have been documented to interact and modulate components of the ECM and alter the determinants of cell adhesion and migration^52^, were also deviantly expressed, failing to reach the levels observed in controls. The teneurins *tenm3* and *tenm4* were underexpressed in tenocytes and the anterior segment mesenchyme (*col9a2*+) respectively. *Tenm3* was found to be underexpressed in tenocytes, which are cells important for building up the ECM and synthesizing and maintaining tendons^53^. *Tenm4*, which is important in the development of neural crest-derived tissues, was underexpressed in anterior segment mesenchyme (*col9a2*+) and periocular mesenchyme cells, a population of neural crest-derived cells that contribute to the development of ocular structures and play an important role in developing the connective tissue around the eye^47^. Importantly, we did not detect changes in other transcription factors known to regulate the expression of ECM components in these transitions, arguing that *lmx1b* directly modulates cell-type specific connective tissue programs.

## Discussion

Analyzing genetic perturbation experiments at single-cell resolution and whole-embryo scale is challenging: any of hundreds of cell types might be impacted, and many effects will be indirect and will manifest at varying scales from the molecular to the anatomic. Here, we describe Hooke and Platt, software tools that work together to statistically model how perturbations impact cells and genes throughout complex tissues or even whole embryos over time. Through differential analysis of many genetic perturbations, the tools organize cell types in an atlas into graphs that describe how cells transit through distinct molecular states. State graphs guide differential expression analysis to identify genes that change as cells differentiate and make fate decisions, helping to prioritize potential perturbation experiments that test whether putative regulatory genes are required for each cell type. When used to interpret perturbation experiments, state graphs help localize the effects of genetic and chemical perturbations within the lineage, distinguishing the most ancestral cell types impacted from their indirectly affected descendants.

Hooke uses Poisson log-normal network models to describe how cell proportions co-vary, how they change over time, and how they are impacted by genetic, chemical, or environmental perturbations. Hooke models track the kinetics of each cell type’s abundance, enabling users to pinpoint when a perturbation interrupts, delays, or blocks each cell type’s development. Hooke accurately detects differences in cell proportion across a wide range of experimental designs, accounts for batch effects and performs contrasts over time, and accommodates complex experiments with multiple perturbations. We also developed Platt, which interprets Hooke models from one or more perturbation experiments to assemble a map of how cell types depend on one another in the lineage. Platt automatically interprets Hooke’s statistical models to track the emergence, expansion, and extinction of transient cell states in a developing embryo and link states through which cells directly transition. Platt also leverages perturbation experiments that interrupt or block state transitions to discriminate between genuine developmental transitions and artifactual links between transcriptional similar cell states. Together, these tools can be used to identify links between healthy and pathological disease states and also define the lineage relationships and genetic requirements in development.

We validated these two tools through an extensive reanalysis of our atlas of perturbed zebrafish embryos, ZSCAPE. Using Platt, we constructed a state transition graph for the developing zebrafish embryo linking cell types to their direct ancestral states, enabling us to define the transcriptional dynamics of numerous fate decisions. The overwhelming majority of cell state transitions were supported by literature or prior knowledge databases and the gene regulation that occurred as these cell types transitioned through states was strongly enriched for genes known to be critical for their development. The state graph also enabled us to systematically catalog the transcription factors regulated at each fate decision, which in turn nominated genetic requirements for each cell type. Both of these analyses were impossible or impractical to perform when we initially collected the ZSCAPE dataset because we lacked scalable, accurate software for doing so.

To demonstrate that our new map of state transitions in the developing embryo is useful for locating the critical genes for specific cell fate decisions, we performed deep single-cell RNA-seq phenotyping on embryos lacking *lmx1b*, a transcription factor frequently upregulated in cell types that contribute to embryonic connective tissue. Differential analysis of the crispants with Hooke and Platt detected numerous cell types lost or blocked in their development, recapitulating expected effects on podocytes,^54^ the midbrain-hindbrain boundary (MHB),^48^ periocular mesenchyme,^47^ and semicircular canal.^46^ We also observed losses in other cell types that expressed *lmx1b* that have not been previously reported, including both the vacuolated cells and the sheath cells of the notochord, as well as a population of tenocyte-like cells originating from the head mesenchyme.

Many of the affected cell types contribute to connective tissues in the embryo and accordingly upregulate collagens as part of their development. Collagens and teneurins were amongst the genes that were underexpressed in these and other cell types in *Δlmx1b* crispants, consistent with the idea that *lmx1b* is important for regulating extracellular matrix production or organization across diverse connective tissue cell types. The *lmx1b* experiment not only demonstrates the potential of Hooke and Platt for finding new genetic requirements, but also for dissecting the regulators of gene expression programs that are shared or repurposed by diverse cell types.

Together, Hooke and Platt constitute powerful new methods in the toolkit for mapping genetic programs that control development and understand how they go awry in disease. They enable the use of controlled perturbations in hypothesis-driven single-cell sequencing experiments to systematically define how cell types depend on one another and on genes. Such experiments are increasingly commonplace, and without scalable tools for interpreting them, data analysis will be ever more rate limiting. Modeling the effects of a perturbation on the proportions and transcriptomes of all cell types in the embryo is challenging because it requires considering effects on all cell types and genes at once. However, as we show here and in Barkan *et al.,* phenotyping at whole-embryo scale and single-cell resolution helps separate primary effects from more downstream consequences, isolating the cell types and gene programs most immediately impacted by a genetic or chemical perturbation. Jointly analyzing cell abundance and transcriptional phenotypes over a state graph often reveals dysregulation in key genes in progenitor cell types that anticipates losses in their descendant fates, which in turn clarifies the mechanisms by which perturbations lead to phenotypes. While it is challenging to extract mechanistic insights from the extraordinarily complex molecular and cellular effects that embryo-scale single-cell experiments capture, the new computational framework contributed by Hooke and Platt helps locate the cell types and transcriptional changes that are most directly affected. More broadly, Hooke and Platt introduce analytic and statistical strategies that become available only in well-controlled, well-powered “atlas-scale” single-cell experiments. We anticipate that new tools will build on or elaborate these strategies in innovative ways to leverage the tremendous potential for embryo-scale single-cell perturbation experiments to reveal parts of the genetic program of vertebrate development.

## Methods

### Benchmarking simulations

Our ability to detect cell proportion changes between different contrasts is a function of the following: abundance of a given cell type, the number of replicates, and the effect size of the perturbation.

To evaluate power, we carried out a simulation analysis:

1. We selected 140 cell types, their abundances ranging from 1 to 20% from a 24 hpf embryo. The proportions of these cell types served as the basis for our simulations.
2. We simulated groups of wild type samples with 5, 10, 15, … 50 replicates in each group. For each replicate, the simulated number of cells of each cell type was calculated as the product of: (a) the cell-type proportions, simulated by fitting a Dirichlet model based on the real proportions from step 1; and (b) the total number of cells recovered for that replicate, simulated on the basis of the mean (n = 1,000) and standard deviation of the cell numbers across replicates in the real dataset.
3. We simulated ten groups of ‘mutant’ samples by repeating the above step except adding shifts to the numbers of cells within each cell type. The shifting scales were based on different effect sizes (0.25, 0.5, 0.75). For instance, effect size = 0.25 represents a 25% reduction in the number of cells.
4. We fit a Hooke model to test whether the cell-type proportions significantly changed between simulated ‘wild type’ and ‘mutant’ samples. We then checked the results stratified by cell type (with different abundances), the number of replicates, and the effect size.

### Comparison with other tools

Hooke was compared to Propeller^14^ (version 0.99.7) and to the method for differential described in Saunders et al, 2023. Briefly, that method fits a generalized linear model with a Beta Binomial response to each cell type’s normalized counts across embryos. Wild type and mutant embryos were simulated as described above. Cell types were mutated in 4 different proportions at a range of effect sizes (.25, 0.5, 0.75) and embryo sizes (250, 500, 1000). These simulations were generated using 100 seeds for each condition to calculate a TPR and FPR.

### scRNA-seq analysis

After RNA and hash-quality filtering, data were processed using the Monocle3 (develop branch, v.1.3.1) workflow using defaults except where specified: estimate_size_factors(), preprocess_cds() with 100 principal components (using all genes) for whole-embryo and 50 principal components for subsets, align_cds(residual_model_formula_str = “∼log10(n.umi)”), and reduce_dimension(max_components = 3, preprocess_method = ‘Aligned’).

### Hierarchical annotation and subclustering

To build maps where cluster annotations corresponded broadly to cell types, the global reference dataset was split into 30 partitions, based on previous tissue annotations. Each partition was re-processed, embedded in three dimensions with UMAP, and subclustered. Clusters were then assigned annotations based on the expression of marker genes (using the top_markers() function) and literature by using the ZFIN database (zfin.org). Each cluster was assigned on ‘cell_type’ annotation. These subtype annotations were manually merged into ‘cell_type_broad’ classifications based on cluster proximity or cell-type functional groupings. Annotations were further merged into ‘tissue’ groups based on whether broad cell types together composed a broader tissue.

### Query dataset projection and label transfer

Projection and label transfer were performed similarly to Saunders *et al*., 2023.

The PCA rotation matrix, batch-correction linear model, and UMAP transformation were computed and saved during the processing of the reference dataset. This computation was done on two levels: first, with all combined reference cells (global reference space), and second, in each of the thirty subgroups (subreference space). One of the projection group labels (e.g. mesoderm, skeletal muscle, hindbrain) was transferred using the majority label of its annotated nearest neighbors (*k* = 10). Nearest neighbors were calculated using annoy, a fast, approximate nearest-neighbor algorithm (https://github.com/spotify/annoy, v.0.0.20). The query dataset was split into 30 subgroups based on these assigned projection group labels. Each query subgroup was projected into the subreference spaces using the corresponding saved PCA, batch correction, and UMAP transformation models using the same projection procedure. Cell type labels were transferred in this subspace using the majority vote of reference neighbors (*k* = 10). Cells without neighbors within a preset distance cutoff (*min_nn_dist* = 1) were removed. Cells that projected onto doublet-labeled clusters were also excluded from the analysis.

### Differential cell abundance testing

After cell type annotation, counts per cell type were summarized per embryo to generate an embryo × cell type matrix. The embryo × cell type matrix was stored as a Hooke cell_count_set object. Counts were compared across genotypes and their paired controls. In a Hooke cell_count_model, two PLN models (v1.2.0) are fit. We refer to these as the full and reduced models. The full model is a PLNmodel fit using genotype, timepoint, and batch as fixed effect covariates and offset as a random effect covariate and are modeled as follows:

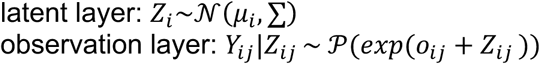

Where Y_ij_ is the observed abundance of cell type *j* in embryo *i* and Z represents the true latent abundances. The μ_i_ parameter corresponds to the fixed effects, and can be decomposed into [inline] where x_i_^T^ is a vector of covariates for each embryo *i* and θ_j_ is a vector of regression coefficients associated to these covariates for cell type *j.* The latent covariance matrix Σ describes the underlying structure of dependence. The fixed quantity o_ij_ is the offset for cell type *j* in embryo *i,* which accounts for expected differences in observed counts due to sampling. For the offset, we calculate an embryo ‘size factor’ by dividing the total number of cells recovered from an embryo by the geometric mean of total cell counts across all embryos.

If the model is fit on wild type only data, the models are fit on timepoint using a natural spline, with an experimental batch term if there are more than two experiments:

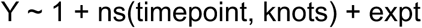

If the model is fit on both wild type and perturbed data, the models are fit using an interaction term between genotype and time with natural splines. A batch term is included if there are more than two experimental batches. Knots in the spline were calculated based on each genotype’s specific collection points.

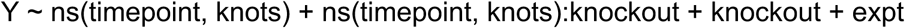

The reduced model is a PLN network model fit and includes nuisance terms such as batch as covariates. It is modeled as follows:

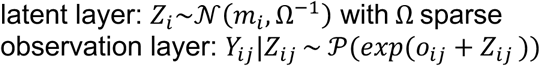

The variance matrix, Σ, captures the correlations between pairs of cell types, while the partial correlations are encoded by its inverse: the precision matrix Ω = Σ^-1^. The precision matrix is assumed to be sparse and a constraint is imposed on coefficients of Ω.

Hooke regularizes the correlations it learns through a penalty scheme that favors transcriptionally similar cell types (e.g. progenitors and their differentiated cell types) based on proximity in transcriptomic (UMAP) space. Cells are first pseudobulked by cell type and the distance between all cell type pairs is then calculated. This resulting distance matrix is contained in a matrix ρ, where d_ij_ is the Euclidean distance between cell types *i* and *j,* and *s* is a scaling constant (by default *s* = 2):

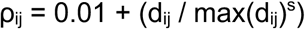

Log abundances are estimated for each condition using the predict() function using the full model, along with standard errors. These abundances are then compared to report a delta log abundance value. P-values were calculated using a Wald test and multiple testing corrected with Benjamini-Hochberg.

### Kinetic modeling

To create the wild type kinetic curves, Hooke models were fit on the wild type data using a natural spline on timepoint, with an experimental batch term if there are more than two experiments. Values are then predicted for a given time interval, with a step size of 2. The same is done for a model fit on mutant data and their corresponding controls, but with perturbation included in the model string. Along with the curves, per-model embryo estimates λ_i_ (i.e. rate parameters, where λ_i_ = exp(Z_i_)) are conditionally predicted given the PLN model, a set of covariates, and the observed normalized cell type counts.

These wild type kinetic curves can additionally be used to estimate the timepoint at which each cell type is at its peak abundance in the reference.

### Lineage construction

Graphs were constructed on each assembly group, first at the cluster-level and then at the cell-type level. For each assembly group, the data is first clustered using Monocle3’s cluster_cells(resolution=1e-3, k=15) to define cell states. A Hooke model is then fit on wild type data with timepoint and experimental batch as covariates. Using this model, the presence of each cell type in a given time interval is determined. Next, a pathfinding graph is initialized over which cells can transition. This starts by computing a partition-based graph abstraction (PAGA) graph,^55^ which represents the connectivity of the UMAP-manifold partitions. Edges with a zero partial correlation in the PLN network model are removed from the PAGA graph. Since the initial direction of flow is unknown, edges are placed in both directions. Next, paths between pairs of states are considered if those states exhibit a reciprocal fold change (one increasing, one decreasing) and have a non-zero partial correlation between them. Paths are then scored by fitting a linear model of:

> time ∼ geodesic distance

The shortest path between these nodes that also aligns with the direction of time is retained in the final graph. Finally, cycles are removed from the graph, resulting in the wild type-only graph.

Next, perturbation data is analyzed. Models are fit for each perturbation in its corresponding experimental time window, with each perturbation model describing a separate experiment that may eliminate one or more cell states. DACTs are considered only if their abundance changes at a timepoint when the corresponding cell type is present in the wild type.

Paths that include a sequence of lost nodes following each perturbation are identified in the pathfinding graph. Paths must also align with the flow of time in control cells (by fitting a linear model), at least over the time window measured in the corresponding perturbation experiment. The same path may recur in multiple perturbation experiments. The level of support for each path is summarized by counting how many edges are supported by perturbation-induced losses. This creates the mutant-supported graph built on cell states.

This process is repeated after contracting the cluster-level graph to cell-type resolution. The graph construction process is repeated on the cell-type, with the contracted graph used as a prior to regularize the partial correlation calculations. For perturbations collected at multiple timepoints, the fold changes are summarized as a weighted average. Each timepoint’s fold change is weighted by the percentage of the maximum wild-type abundance observed at that timepoint. Finally, genetic requirements are calculated where if a cell type is significantly lost and its parent remains unchanged the change is labeled as “direct”; otherwise, it is considered “indirect.”

### Assessing support for Platt graph state transitions

To more systematically assess the accuracy of the map of the lineage transitions assembled by Platt, we scored each edge by the strength of support in our own perturbation data. Directed edges (*u*, *v*) from cell state *u* to state *v* were deemed directly supported by a perturbation when all of the following criteria were true: (1) at least one of them expresses the gene(s) targeted in the perturbation; (2) whenever *u* is a DACT, *v* is significantly reduced; (3) any parent states of *u* are unaffected. If the first two criteria are true, but the last is not owing to changes upstream of *u*, the edge is scored as indirectly supported by the perturbation.

### Aligning cell type labels with previously published ontologies

Each cell type in the atlas was assigned a Cell Ontology identifier (CL id) from ZFIN. If the cell type was at finer resolution than listed on ZFIN, the CL id of its “cell type broad” was used (e.g. cranial motor neuron progenitor becomes cranial motor neuron). Associated anatomical phenotypes from ZFIN were also mapped to the atlas cell types.

### Defining genetic requirements of cell types with Monarch

The Monarch Initiative Database, which integrates data about genes and phenotypes across species, was referenced to define the set of known genetic requirements for each cell type. First, each cell type’s associated CL id was queried in the Monarch database. Then, associated biological process genes and phenotypes were collected into a table. This process is repeated for a cell type’s anatomical id. Finally, each gene is then categorized as direct (a cell type is lost when the gene is affected, but not its parent) or indirect (a cell type and its parent are lost when the gene is affected) using the Platt graph.

### Differential gene expression analysis

Expression values were first aggregated for each embryo across cell types into ‘pseudo-cells.’ Differential gene expression across perturbations was computed for every cell type using a Platt method that models each gene’s expression in each cell type and treatment condition via a generalized linear regression model (GLM) as described previously^20,56^ using the fit_models function in Monocle3 (v1.3.1)^57^.

To prevent inflated effect estimates, such as those from low-expression genes in rare cell types, an empirical Bayes shrinkage approach, ashr^58^, was applied to regularize effect estimates. Log fold changes for inaccurate estimates tend to be shrunken toward zero, while accurately estimated effects, where the standard error is small relative to the estimate, remain unchanged. Unless otherwise specified, shrunken estimates are reported when presenting log-fold changes for individual genes within specific cell types.

### Differential gene expression over reference transitions

Differential gene expression was computed across the Platt cell state transition graph on wild type data. If the Platt graph contains a directed edge from [parent] cell type *x* to [child] cell type *y*, the differential expression test would be the genes differentially expressed in cell type *y* when compared to cell type *x.* These results were classified into different gene expression patterns such as activated, deactivated, upregulated, downregulated, or maintained. Patterns are defined as follows:

- Activated: expressed in self, but not in parent
- Deactivated: not expressed in self, expressed in parent
- Upregulated: expressed in self, expressed in parent, higher than parent
- Downregulated: expressed in self, expressed in parent, lower than parent
- Maintained: expressed in self, expressed in parents, same as parent
- Absent: not expressed in self or parent

If there is a branch point in the graph, genes are additionally labeled with a prefix:

- Specifically: pattern is present in only one of the child cell types
- Selectively: pattern is present in two or more of the child cell types but not all

### Deviantly expressed gene expression analysis

Reference gene expression patterns were compared to the perturbation DEG results to classify genes as “deviantly expressed” (DvEG). Genes are DvEG in each perturbation if they are 1) upregulated during a wild type transition but are underexpressed in perturbed cells undergoing that same transition, 2) downregulated during a wild type transition but are overexpressed in perturbed cells undergoing that same transition or 3) maintained in the wild type transition, but differentially expressed in perturbed cells undergoing the same transition.

### Animal rearing, staging, and stocks

Staging followed (Kimmel *et al.*, 1995) and fish were maintained at 28.5°C under 14:10 light:dark cycles. Fish stocks used were wild type WIK/AB. Fish were anesthetized prior to imaging or dissociation with MS222 and euthanized by overdose of MS222. All procedures involving live animals followed federal, state and local guidelines for humane treatment and protocols approved by Institutional Animal Care and Use Committees (protocol #4405-02) of the University of Washington.

### In situ hybridization, immunohistochemistry and labeling

Colorimetric in situ hybridization used digoxigenin labeled probes using standard conditions (Thisse & Thisse, 2007). Tyramide signal amplification was performed according to the protocol by Lauter *et al*., (2011).

### Imaging

Embryos live-imaged were anesthetized with MS222 and photographed on a Nikon AZ100 microscope. Embryos were put into 70% glycerol and imaged on a Nikon AZ100 microscope. Images were corrected for color balance and display levels as necessary with all conditions within each analysis grouping corrected identically.

### CRISPR-Cas9 mutagenesis in zebrafish embryos

gRNAs were designed using either the Integrated DNA Technologies (IDT) or CRISPR online tools. gRNA and RNP preparation closely follow a recently published protocol for efficient CRISPR–Cas9 mutagenesis in zebrafish^20^. Briefly, gRNAs were synthesized as crispr RNAs (crRNAs, IDT), and a 50 µmol crRNA:trans-activating crispr RNA (tracrRNA) duplex was generated by mixing equal parts of 100 µmol stocks. Cas9 protein (Alt-R S.p. Cas9 nuclease, v.3, IDT) was diluted to a 25 µmol stock solution in 20 nmol HEPES-NaOH (pH 7.5), 350 mmol KCl, 20% glycerol. The RNP complex mixture was prepared fresh for each injection by combining 1 µl 25 µmol crRNA:tracrRNA duplex (with equal parts each gRNA per gene target), 1 µl of 25 µmol Cas9 Protein and 3 µl nuclease-free water. Before injection, the RNP complex solution was incubated for 5 min at 37 °C and then kept at room temperature. Approximately 1–2 nl was injected into the cytoplasm of one-cell-stage embryos.

### Chemical Inhibitor Screen

Embryos were exposed to the inhibitors 100µM Cyclopamine (Cayman, cat. no. 11321) at shield stage with vehicle controls 1% / 239 µM Ethanol for Cyclopamine control. Embryos were exposed to the inhibitors 20µM WntC59 (Cayman, cat. no. 16644) at 8-somite stage with vehicle control 0.24% / 33.8 µM DMSO. Embryo media was replaced and inhibitors or vehicle were replenished every 24 hours until time of collection.

### Preparation of Barcoded Nuclei

Individual zebrafish embryos (18, 24, 36, 48 and 72 hpf) were manually dechorionated with forceps and transferred to a 10cm petri dish containing 1X TrypLE (Thermo Fisher, cat. no. 12604013) and MS222 (Millipore Sigma, cat. no. 886-86-2). Embryos were dissociated into single cells following the protocol described in Saunders *et al.*, 2023. Cell lysis and fixation followed the protocol described in Martin *et al.*, 2023, with an additional 5μl of C6 amine-modified hash DNA oligo (10uM, IDT, 5′-/5AmMC12/GTCTCGTGGGCTCGGAGATGTGTATAAGAGACAG[10bp barcode]BAAAAAAAAAAAAAAAAAAAAAAAAAAAAAAAA -3’) mixed into the Hypotonic Lysis Buffer Solution B.

### sci-RNA-seq3 library construction

The fixed and hashed nuclei were processed according to the following protocol https://www.ncbi.nlm.nih.gov/pmc/articles/PMC9839601/pdf/nihms-1846803.pdf (Martin *et al.*, 2023)

### Sequencing, read processing, and cell filtering

Sequencing, read processing, and cell filtering were performed according to Saunders *et al.*, 2023. An enrichment cutoff of 2.5 was set based on the distribution of enrichment ratios.

## Data availability

Processed data files used in the paper analysis are available for download at https://cole-trapnell-lab.github.io/lmx1b/ under CC-BY-NC. The accession number for the scRNA-seq data reported will be available soon.

## Code availability

Pipelines for generating count matrices from sci-RNA-seq3 sequencing data are available at https://github.com/bbi-lab/bbi-dmux and https://github.com/bbi-lab/bbi-sci. Analyses of the single-cell transcriptome data were performed using Monocle3, Hooke, and Platt; general tutorials can be found at https://cole-trapnell-lab.github.io/monocle3/, https://cole-trapnell-lab.github.io/hooke/, and https://cole-trapnell-lab.github.io/platt/.

## Project Acknowledgements

We thank the Brotman Baty Institute Advanced Technology Lab for support with sequencing and the data processing pipeline. We thank Dr. Dave Raible, Dr. Ali Shojaie, Dr. Robert Waterston, and Dr. Jay Shendure for helpful discussions as we were developing this project.

## Author Contributions

M.D. and C.T. conceived the project. E.B., A.T., H.L., N.L., did dissociation, nuclei collections, and sci-Plex experiments. M.D., A.T., E.B., M.C., R.Z.F., D.K., and C.T. annotated cell types in the reference. M.D., C.T., B.E. and R.Z.F. wrote and maintained the software. M.D. performed computational analyses. D.K. performed the WISH. M.D. and C.T. wrote the manuscript with input from all co-authors. C.T. supervised this project.

## Corresponding Authors

Correspondence to Cole Trapnell.

## Supplemental Information

**Supplemental Table 1** - Genes used for cell type annotation

**Supplemental Table 2** - DACTs by gene target and timepoint

**Supplemental Table 3** - Edges support by literature

**Supplemental Figure 1.**
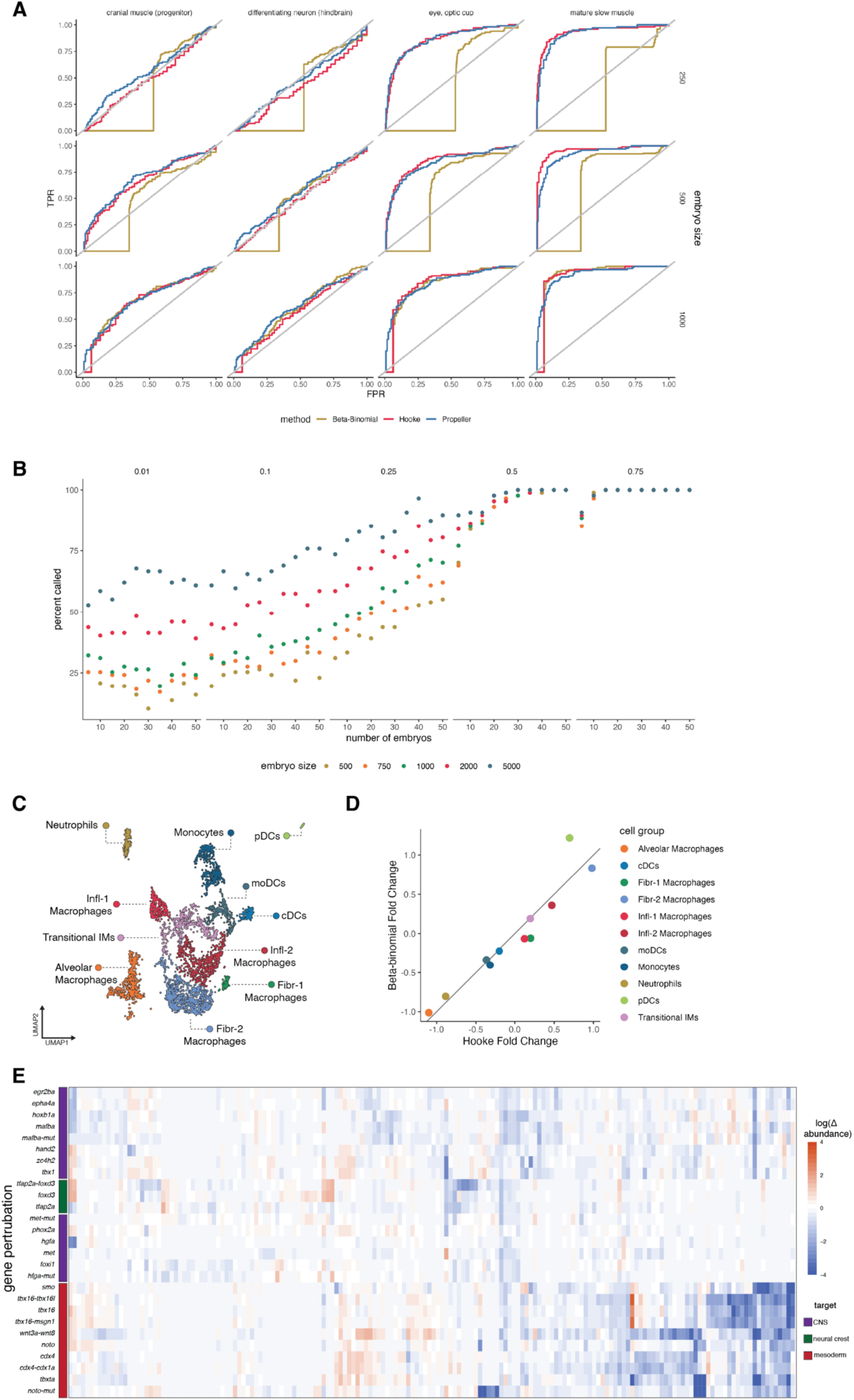
Evaluating the performance of Hooke. A) Comparison of Hooke to other differential abundance tools. Receiver operating characteristic (ROC) curves comparing the true positive rate (TPR) and false positive rate (FPR) of Hooke to Beta-Binomial and Propeller. B) Evaluation of Hooke’s power to detect a range of effect sizes across embryo size and number of embryos. C) UMAP of myeloid cells colored by cell type. D) Hooke effect sizes compared to Beta-Binomial effect sizes. E) Summary of fold change effects of 23 genetic perturbations across the time range for each cell type in the embryo dataset. “Gene”-mut refers to null mutants rather than crispants.

**Supplemental Figure 2.**
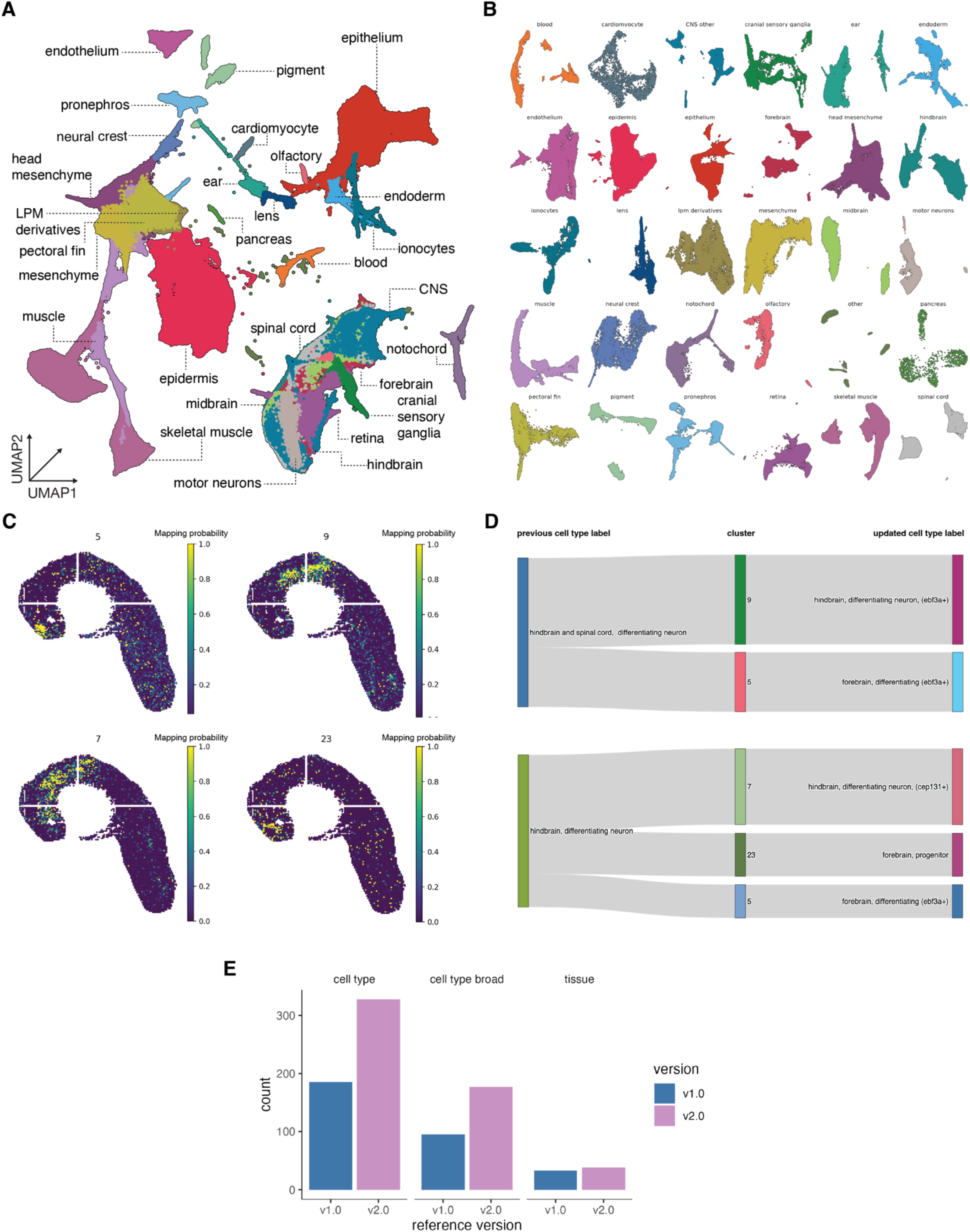
Reference v2.0. A) UMAP of reference v2.0 plotted in global space colored by tissue. B. Subspace UMAPs of each tissue. C) Spatial origin of previously labeled differentiating hindbrain clusters inferred from the Liu dataset and Tangram Algorithm. D) Sankey plots mapping previous cell type labels to new spatial labels. E) Comparison of Reference 1.0 vs Reference 2.0. Bar plots compare the counts of reference annotations at three resolutions: cell type, cell type broad, and tissue

**Supplemental Figure 3.**
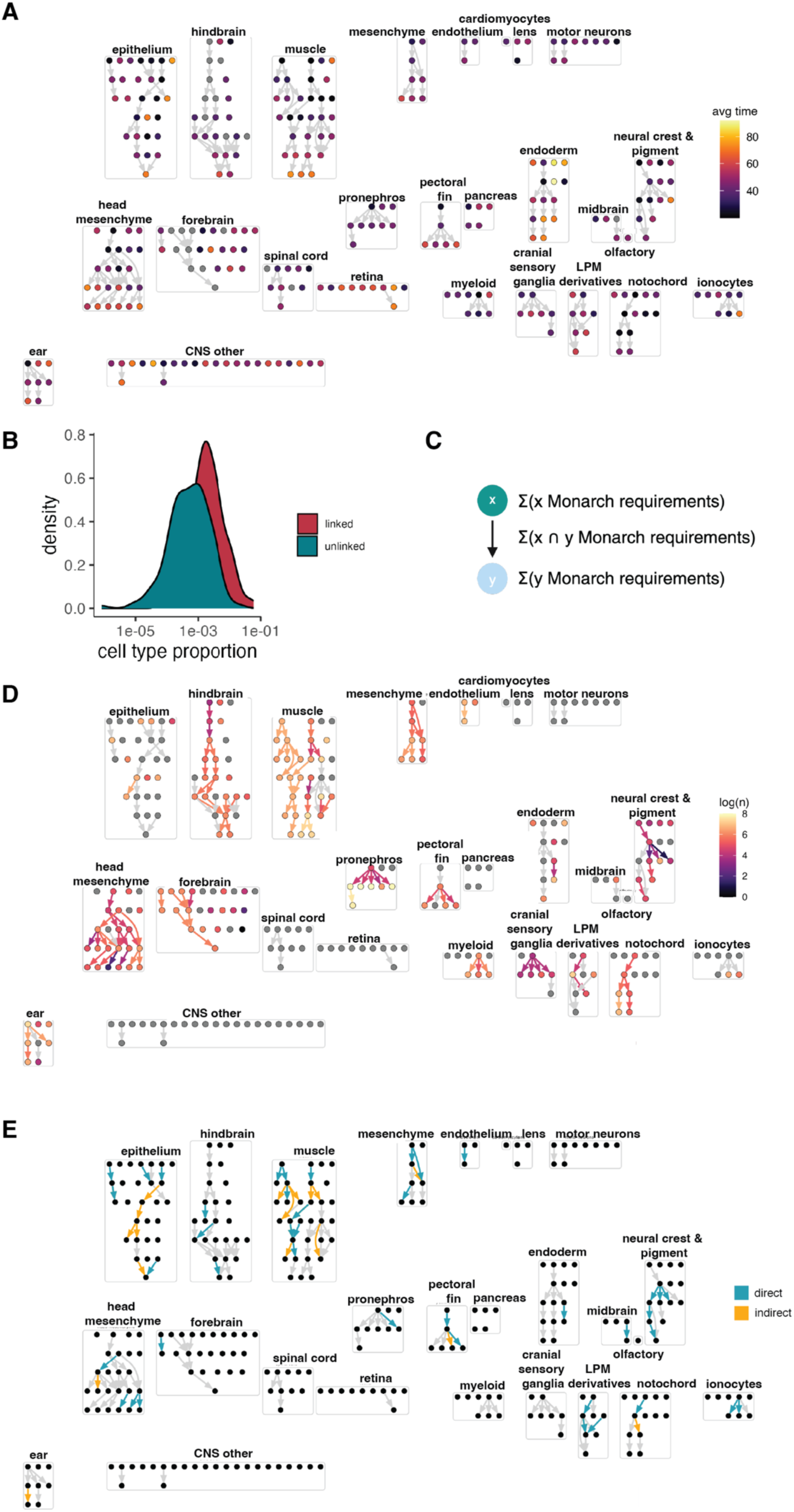
Evaluating Platt graph support. A) Platt graph colored by average timepoint of each cell type. B) Density plot comparing the cell type proportion of unlinked vs linked cell types in the Platt graph. C) Schematic explanation of calculating monarch edge support. D) Platt graph with nodes and edges colored by amount of monarch support. E) Platt graph colored by edges with direct and indirect support from ZSCAPE perturbation data.

**Supplemental Figure 4.**
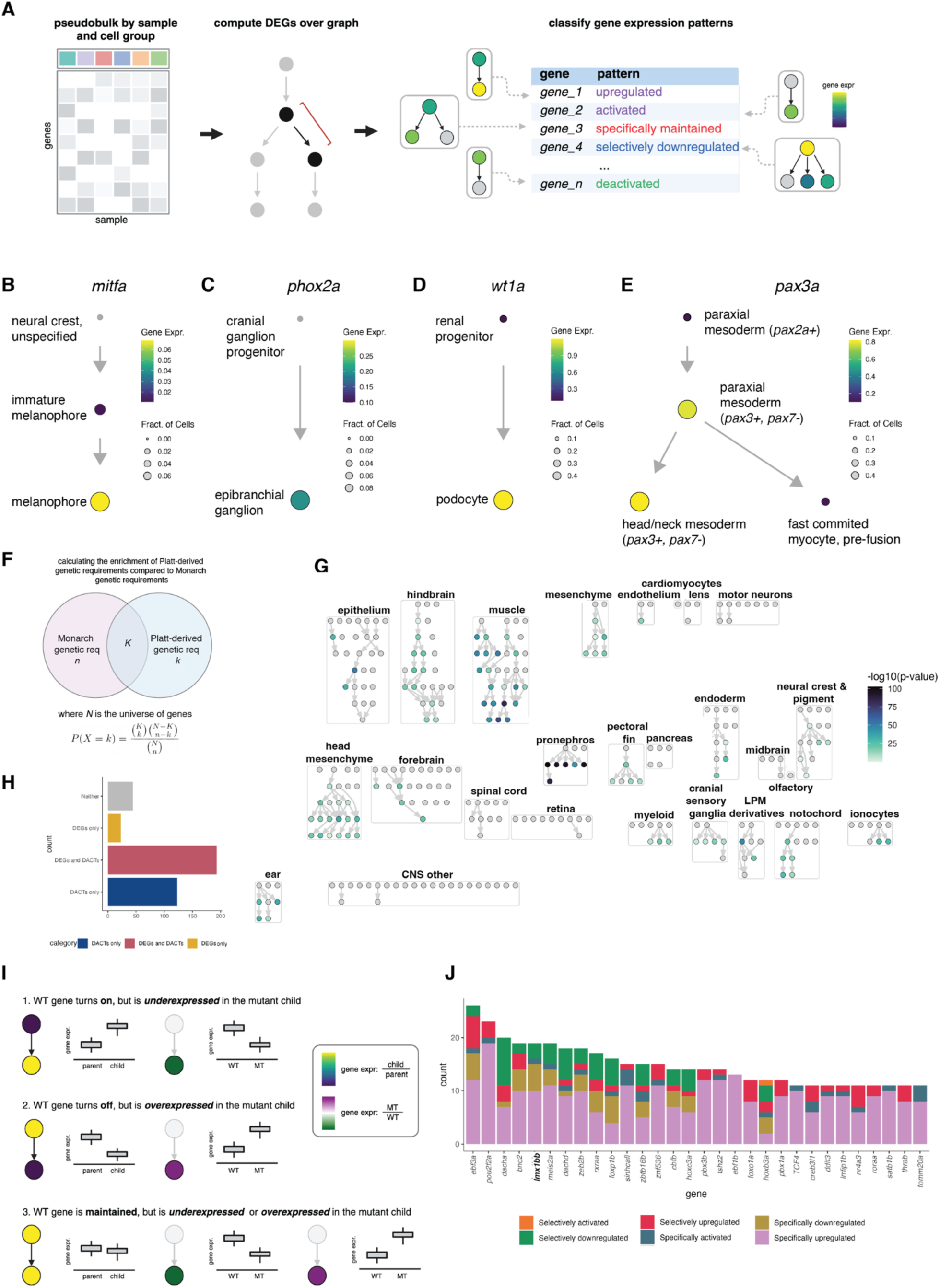
Calculating DEGs across the graph. A) A schematic of how Platt identifies differentially expressed genes. It uses an inferred or provided state transition graph to guide the differential expression testing and classifies these genes into different pattern types. Activated: expressed in self, but not in parent; Deactivated: not expressed in self, expressed in parent, no siblings; Upregulated: expressed in self, expressed higher in parent; Downregulated: expressed in self, expressed lower in parent; Maintained: expressed in self and in parents. B-E) Subsection of Platt graphs colored by gene expression of genes that were identified as active, upregulated, or selectively upregulated. F) Schematic of the calculation of the enrichment of Platt-derived genetic requirements compared to Monarch-derived genetic requirements. G) Platt graph colored by the -log_10_(p-value) of a hypergeometric test between Platt-derived genetic requirements and Monarch-derived genetic requirements. H) Bar plot comparing counts of nodes with DACTs, DEGs, both or none. I) Schematic representation of how DvEGs are determined. J) Barplot of the number of times each transcription factor has a differentially expressed pattern across lineages. Colors refer to pattern interpretation categories.

**Supplemental Figure 5.**
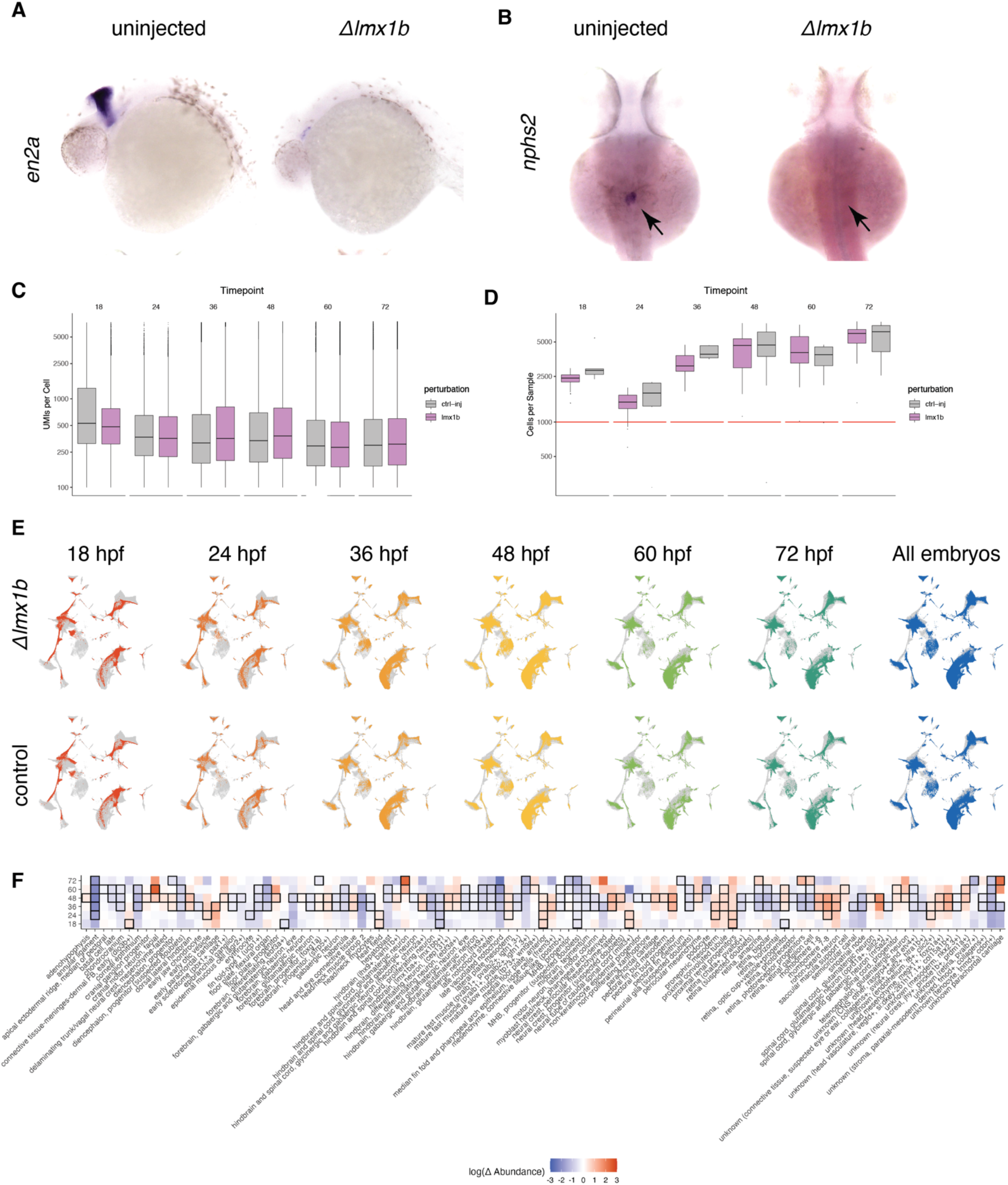
Experimental metrics for *Δlmx1b* experiment. A) In situs of *Δlmx1b* embryos, *en2a* is a marker of the midbrain-hindbrain boundary B) and *nphs2* is a marker of podocytes. C) Distribution of the highest UMI hash barcode per cell divided by the second highest UMI hash barcode per cell. Cells with a ratio of 2.5 or greater, after subtracting background hash UMI level, were included in our study. D) Boxplot of the number of UMIs per cell across all embryos per perturbation (color) and timepoint. E) boxplot of the number of cells per embryo collected across all embryos per perturbation (color) and timepoint. F) UMAPS of *Δlmx1b* and control cells across timepoints. G) Heatmap of fold change in abundance in all cell types for each *Δlmx1b* timepoint collection vs controls. Black boxes denote significance (q<0.05, PLN regression).

**Supplemental Figure 6.**
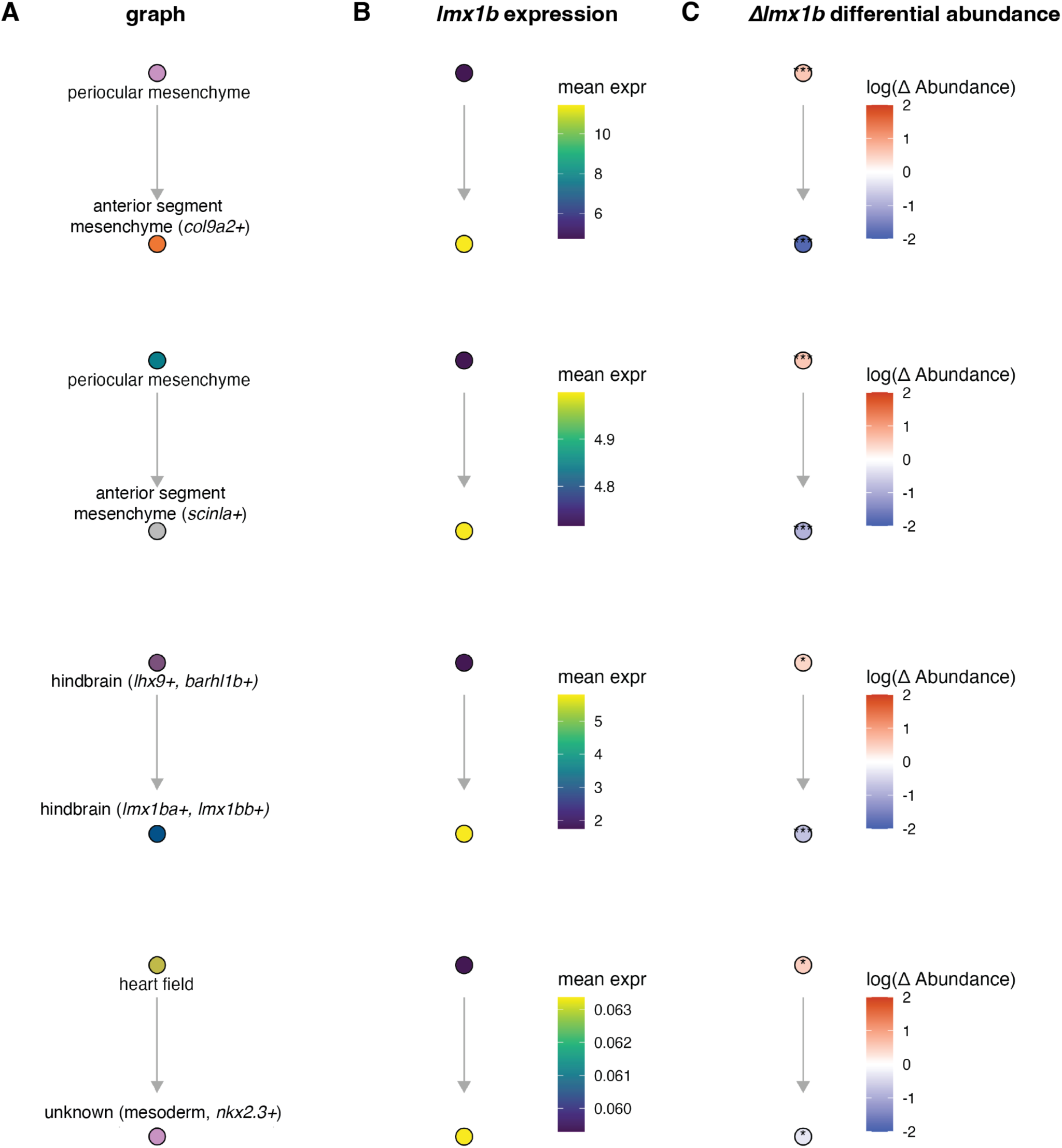
Edges supported by the *Δlmx1b* crispant. A) Selected edges supported by *lmx1b*. B) Platt graphs colored by Hooke abundance change in *Δlmx1b* crispant. C) Platt graphs colored by the sum of *lmx1ba* and *lmx1bb* wild type gene expression.

**Supplemental Figure 7.**
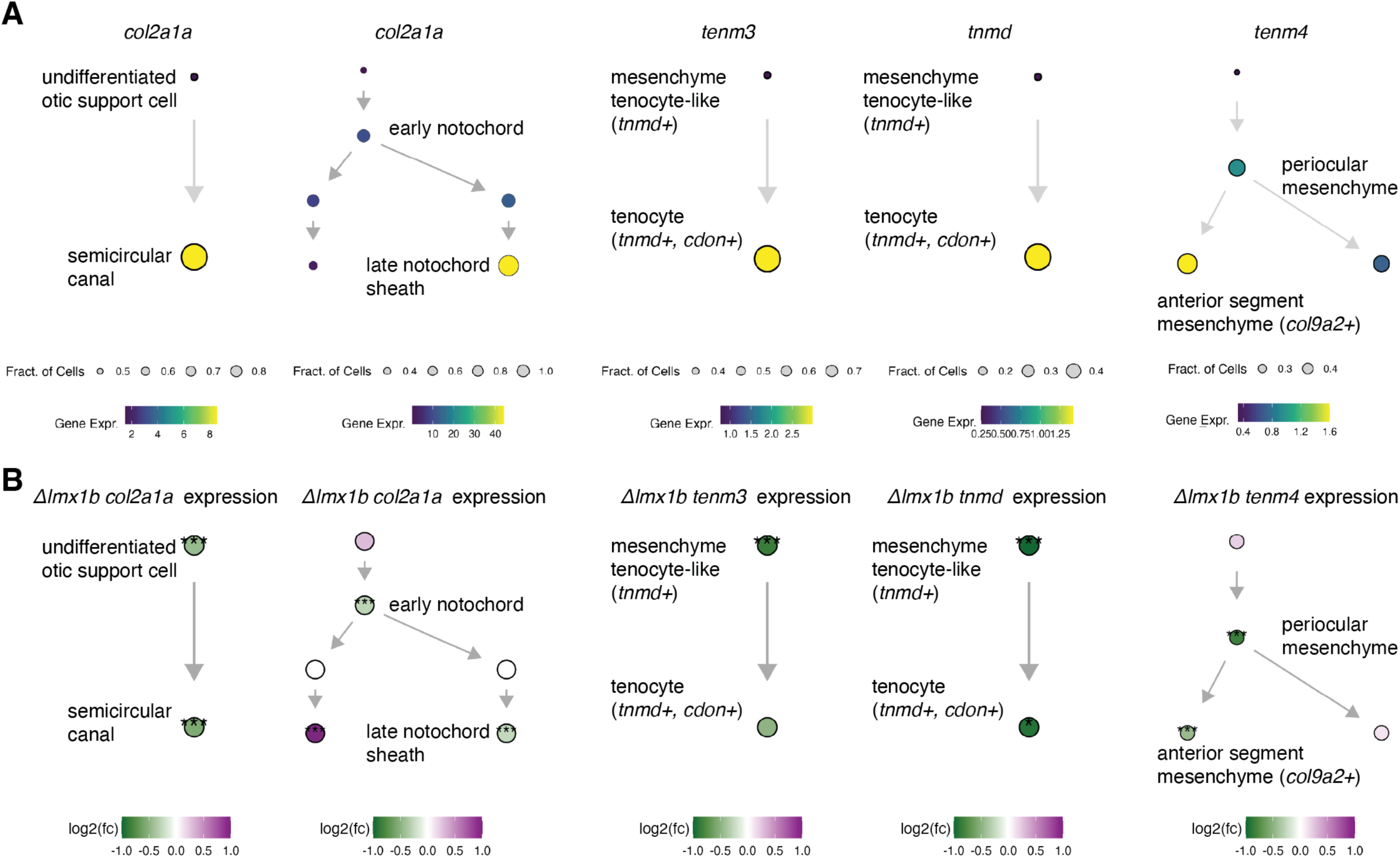
Deviant *Δlmx1b* DEGs. A) Platt state graphs colored by wild type gene expression of select genes that turn on in the graph section. B) Platt state graphs colored by the log fold change of select genes in *Δlmx1b* compared to wild type (*: p < 0.05, **: p < 0.01, ***: p < 0.001).

